# Highly structured homolog pairing reflects functional organization of the *Drosophila* genome

**DOI:** 10.1101/443887

**Authors:** Jumana AlHaj Abed, Jelena Erceg, Anton Goloborodko, Son C. Nguyen, Ruth B. McCole, Wren Saylor, Geoffrey Fudenberg, Bryan R. Lajoie, Job Dekker, Leonid A. Mirny, Ting (C.-ting) Wu

## Abstract

*Trans*-homolog interactions encompass potent regulatory functions, which have been studied extensively in *Drosophila,* where homologs are paired in somatic cells and pairing-dependent gene regulation, or transvection, is well-documented. Nevertheless, the structure of pairing and whether its functional impact is genome-wide have eluded analysis. Accordingly, we generated a diploid cell line from divergent parents and applied haplotype-resolved Hi-C, discovering that homologs pair relatively precisely genome-wide in addition to establishing *trans*-homolog domains and compartments. We also elucidated the structure of pairing with unprecedented detail, documenting significant variation across the genome. In particular, we characterized two forms: tight pairing, consisting of contiguous small domains, and loose pairing, consisting of single larger domains. Strikingly, active genomic regions (A-type compartments, active chromatin, expressed genes) correlated with tight pairing, suggesting that pairing has a functional role genome-wide. Finally, using RNAi and haplotype-resolved Hi-C, we show that disruption of pairing-promoting factors results in global changes in pairing.

**One Sentence Summary:** Haplotype-resolved Hi-C reveals structures of homolog pairing and global implications for gene activity in hybrid PnM cells.

## Main Text

Major hallmarks of chromatin organization include chromosome territories (*1*), compartments of active (A-type) and inactive (B-type) chromatin as delineated by the conformation capture technology of Hi-C, as well as chromosomal domains variably known as contact domains or topologically associating domains (*2-7*). These layers of organization encompass countless *cis* and *trans* interactions that determine the 4D organization of the genome. Among *trans* interactions, an important class includes those occurring between homologous (*trans*-homolog) as versus heterologous (*trans*-heterolog) chromosomes. Although long considered relevant only to meiosis, *trans*-homolog interactions are now widely recognized for their capacity to affect gene function (reviewed by (*8-12*)). What remains unclear is the global impact of such interactions. In this study, we examine the first genome-wide maps of *trans*-homolog interactions in a newly established hybrid *Drosophila* cell line and then assess the relationship of those interactions to genome function.

Our studies take advantage of *Drosophila*, where *trans-homolog* interactions are especially well-studied. Here, homologs are paired in somatic cells throughout nearly all of development (*8*). In fact, *Drosophila* is the first organism in which *trans*-homolog interactions were implicated in gene regulation, as it is here that somatic pairing was discovered (*13*) and subsequently found to impact intragenic complementation (*14*). These and other foundational studies documenting the impact of *trans*-homolog interactions on genome function have relied heavily on genetic approaches to infer pairing, with recent studies also making use of fluorescent *in situ* hybridization (FISH) to visualize pairing directly ((*8-12, 15, 16*); see (*17*) for live imaging). In addition, recently, juxtaposed chromosomal domains have been detected by super-resolution imaging (*16, 18*). Somatic pairing has now been implicated in numerous biological phenomena across a diversity of species, including mammals, with pairing-dependent gene regulation, a well-recognized form of transvection, being among the best understood (reviewed by (*8-12*)). In *Drosophila*, transvection has been observed at many loci, suggesting that pairing may even function as a regulatory mechanism genome-wide (*19-22*). Recently, this view has been supported by computational simulations of homolog pairing in *Drosophila (23)*. Thus, the question as to whether pairing can serve as a genome-wide regulatory mechanism is drawing increasing attention.

Our work used haplotype-resolved Hi-C to examine the detailed architecture of homolog pairing in diploid *Drosophila* cells and its potential genome-wide impact on gene regulation. Haplotype-resolved Hi-C has been used to investigate *cis* interactions within mammalian genomes (*24-35*) and diploid homolog pairing in yeast (*36*) and, in our companion paper (Erceg, AlHaj Abed, Golobordko *et al. bioRxiv* (*37*)), we developed a methodology for applying this approach to study homolog pairing in hybrid *Drosophila* embryos. In that study, we demonstrated pairing to be genome-wide and provided a framework in which to consider pairing in terms of precision, proximity, and continuity. We further revealed *trans*-homolog derived domains that are positionally concordant with domains defined by *cis* interactions as well as uncovered a potential connection between pairing and chromatin accessibility in early embryogenesis.

In the current study, we generated a diploid cell line from hybrid *Drosophila* embryos, produced from crossing divergent parents. Taking advantage of the greater homogeneity of cell lines and higher pairing levels, we applied haplotype resolved Hi-C to achieve a high-resolution map of homolog pairing, observing telomere-to-telomere alignment of homologs and revealing *trans*-homolog domains and compartments. In addition, we detailed the variation in the structure of pairing, documenting an extensive interspersion of tightly paired regions with loosely paired regions across the genome. Excitingly, we also found a strong association between pairing and active chromatin, compartments, and gene expression, thus resolving the long-standing question of whether pairing can bear a genome-wide relationship to gene expression.

We began our study by crossing two strains of the *Drosophila* Genetic Reference Panel lines (057 and 439) that differ by ~5 single nucleotide variants (SNVs) per kilobase (kb) (table S1) (Erceg, AlHaj Abed, Golobordko *et al. bioRxiv (37)*) to generate 2-14 hour old embryos that were homogenized, spontaneously immortalized, and then serially diluted to generate clonal cell lines (Fig. 1A; Supplementary Material). The clonal line used in this study, Pat and Mat (PnM), homogeneously expresses myocyte enhancer factor 2, suggesting it to be of mesodermal origin (fig. S1A, B). Karyotyping, in combination with homolog-specific fluorescent *in situ* hybridization (FISH) proved PnM cells to be male, diploid, and hybrid, with only chromosome 4 showing irregularities (Fig. 1B, C; Supplementary Material). Finally, FISH analyses targeting two heterochromatic and three euchromatic loci confirmed high levels of pairing (Fig. 1D).

**Fig. 1.**
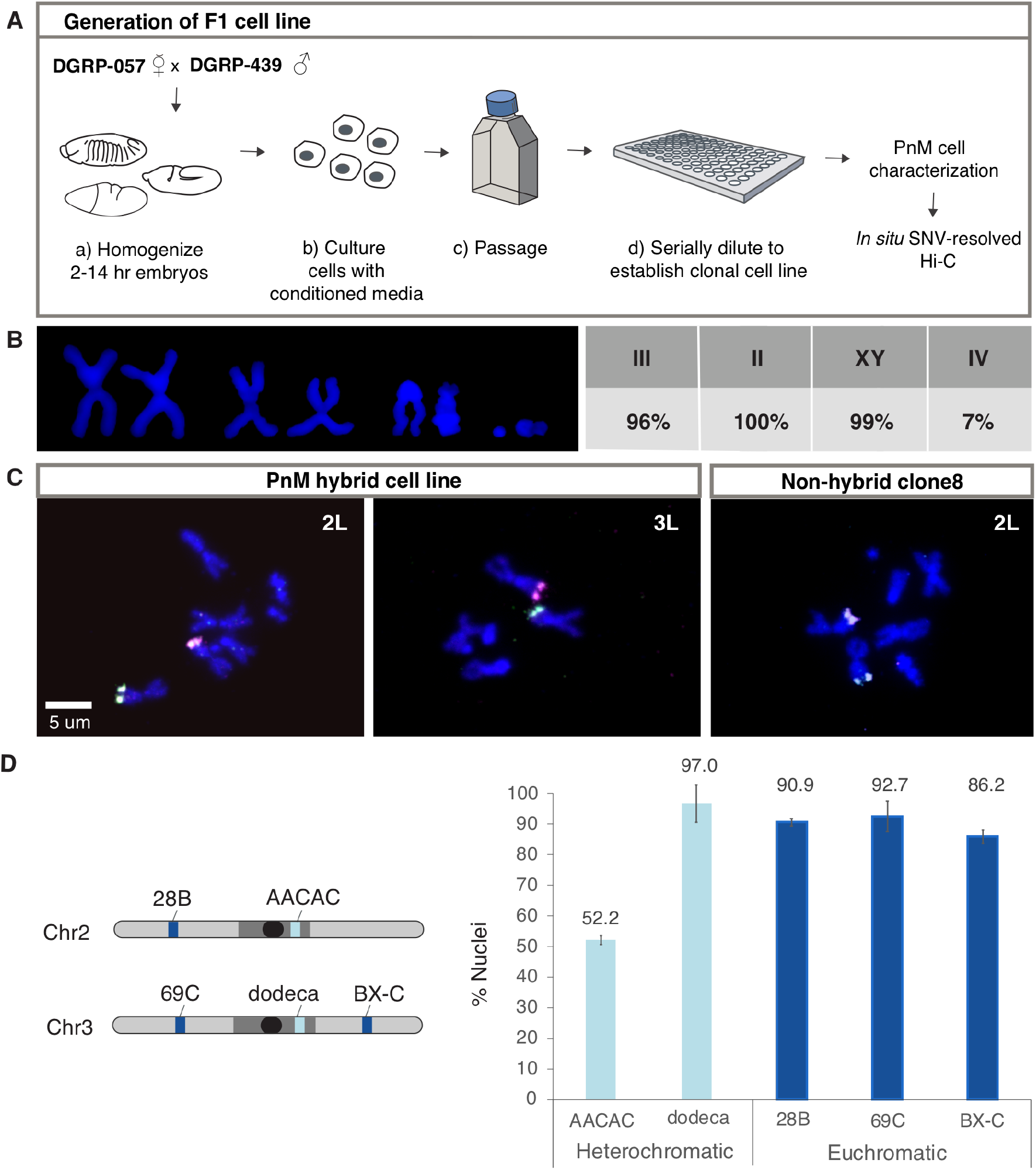
PnM cell line. (A) Generation of the cell line. (B) Karyotyping demonstrates PnM cells to be male and diploid (N = 50). (C) HOPs probes distinguishing 057-derived (magenta) from 439-drived (green) homologs for chromosomes (Chr) 2 and 3 on metaphase spreads for PnM and control (non-hybrid) clone 8 cells confirmed PnM to be hybrid. (D) Left: Locations of heterochromatic (blue) and euchromatic (grey) chromosomal regions targeted by FISH. Oval, centromere. Right: Levels of pairing in PnM cells quantified as percent of nuclei in which FISH signals, representing allelic regions, co-localized (center-to-center distance between signals ≤0.8 μm; error bars, s.d. for two biological replicates; N > 100 nuclei/replicate).

We next performed haplotype-resolved Hi-C on PnM cells. In this form of Hi-C, each of the two fragments of genomic DNA that are ligated together by virtue of their proximity *in situ* are assigned a parental origin based on the SNVs they carry, thus permitting researchers to distinguish Hi-C read pairs that represent *cis* maternal*, cis* paternal*, trans*-homolog (*thom*), and *trans-heterolog* (*thet*) interactions. By requiring at least one SNV per side of each read pair, we obtained 75.4 million mappable read pairs, producing a 4 kb resolution haplotype-resolved map of the mappable portion of the genome (e.g., excluding repetitive regions), wherein less than 0.4% of *thom* read pairs are expected to have resulted from read misassignment (Supplementary Materials; fig. S2A, table S2).

As shown in Figure 2A, homologs are aligned genome-wide, comparable to the global *thom* signature detected in early *Drosophila* embryos (Erceg, AlHaj Abed, Golobordko *et al. bioRxiv (37)*). Strikingly, however, *thom* read pairs were ~7.8 times more abundant in PnM cells than in *Drosophila* embryos (fig. S2B). In addition, when considering *thom* contacts as a function of the separation of loci along the genome (genomic separation), we found them to be more abundant at all genomic separations (fig. S2C). These observations agree with the higher levels of pairing observed by FISH in PnM cells as compared to developing embryos (Fig. 1D), due possibly to a greater percentage of cells with paired homologs, an increased fraction of the genome exhibiting pairing, and/or a smaller proportion of dividing cells in the PnM cell line (fig. S6). Importantly, the greater abundance of *thom* contacts argued that an analysis of pairing in PnM cells would yield new insights into the structure of paired homologs.

**Fig. 2.**
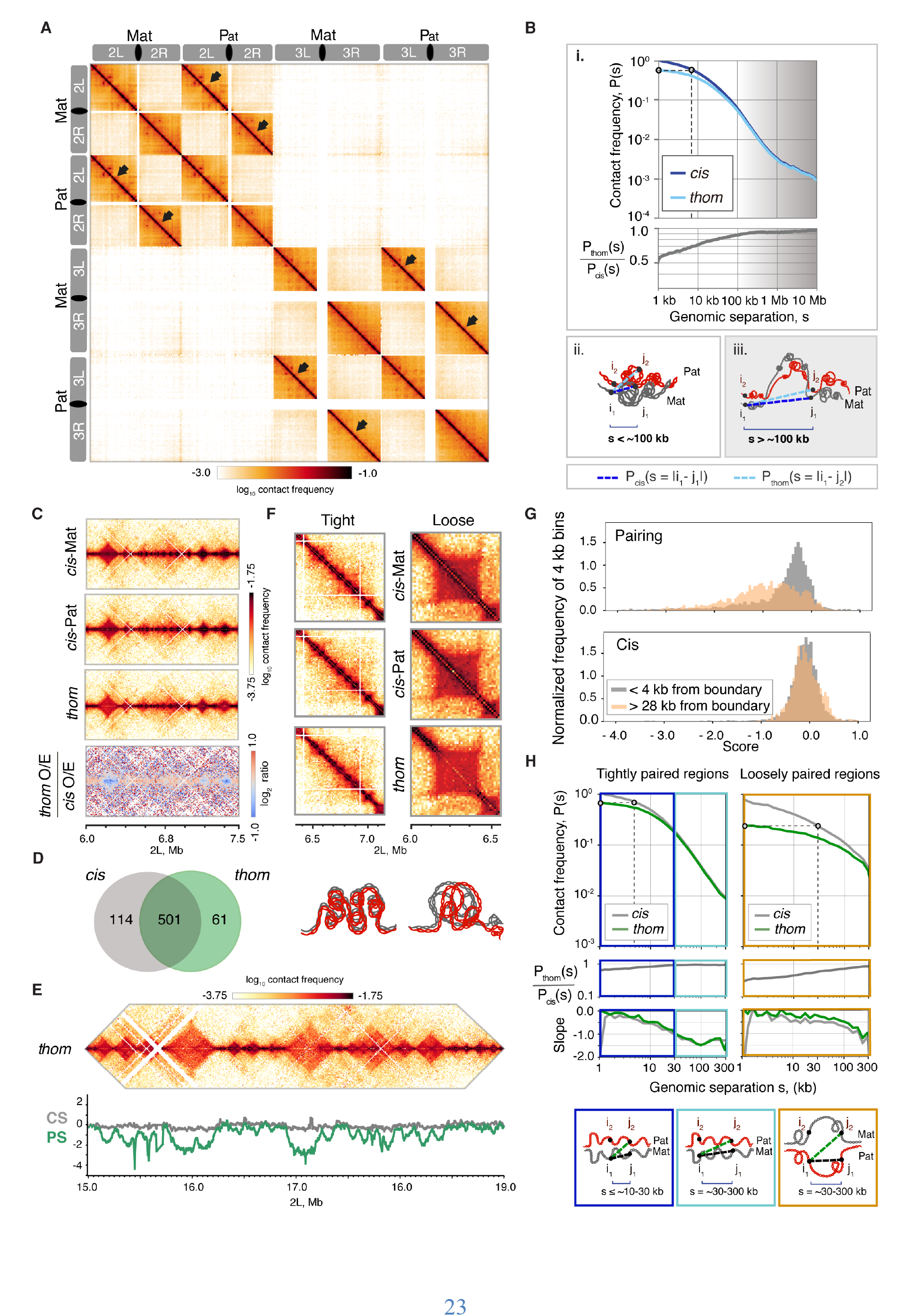
Haplotype-resolved Hi-C detected genome-wide in-register pairing, as well as *thom*-domains and variation in the structure and precision of pairing. (A) Hi-C contact map for the left (L) and right (R) arms of Chr2 and 3. (Bi) Top, *cis* and *thom* contact frequencies, P(s), plotted against genomic separation, s, normalized to the *cis* frequency at s = 1 kb. Dotted line: *thom* contacts at s = 1 were as frequent as *cis* contacts at s = 8 kb. Bottom, ratio of *thom* to *cis* contact frequencies. Note differences in c*is* and *thom* contact frequency at separations, s < ~100. At s > ~100 kb, shaded region, *thom* contact frequency decay was concordant to *cis*. Bii-iii. Dashed lines indicate that the two loci that are close enough to each other to be captured by Hi-C, and represent P_thom_(s) as a function of separation s = |i_1_-j_2_| for loci i_1_ and j_2_ located on different homologs, compared to P_cis_(s) for loci i_1_ and j_1_ separated by distance s = |i_1_-j_1_| on the same chromosome. P(s) at s < ~100 kb, left panel. P(s) at s > ~100 kb, right panel, shaded region in (B)i.(C) *Cis* and *thom* contact maps for a 1.5 Mb region on 2L were highly concordant with each other (top two panels), and with the *thom* map (third panel). *Thom/cis* map (bottom panel) showed a high *thom* concordance with *cis*, apart from a depletion in contacts in some domains, in blue. (D) Overlap of domain boundaries as defined by *cis* and *thom* contacts. (E) Top, *thom* domains, and insulating boundaries within a 4 Mb region of 2L. Bottom, pairing score (PS, green) and *cis* score (CS, grey). (F) Examples of tight and loose pairing, with corresponding schematics of possible structures. (G) Distributions of PS and CS near a boundary (less than 4 kb) in grey, or far from boundary (more than 28 kb away), in orange, showed higher pairing near boundaries. (H) Top, *cis* and *thom* contact frequencies within tight and loose regions (normalized as in Figure 2B) plotted against genomic separation. Middle, ratio of *thom* to *cis* contact frequencies, bottom, shows slopes. Tightly paired regions represented two modes of decay, shallow (dark blue) and steep (light blue), while loosely paired regions showed one mode, which was shallow (orange). Dotted lines: *thom* contacts at s = 1 were as frequent as *cis* contacts at s = ~5 kb and ~30 kb in tightly and loosely paired region, respectively. Schematics illustrate how the differences in P_thom_(s) and P_cis_(s) between tight and loosely paired regions may reflect differences in their organization.

We began by comparing the probability with which a locus will interact with another locus in *cis* as versus in *thom* at varying genomic separations. We reasoned that, if pairing were maximally precise, tight, and continuous (‘railroad track’), any two loci would interact in *thom* nearly as often as they interact in *cis* regardless of genomic separation. In contrast, in the case of imprecise, loose, discontinuous pairing, the relative frequencies of *cis* and *thom* contacts could differ quite substantially. Note that this railroad track pairing does not preclude long-range *thom* interactions as paired chromosomes can fold back onto themselves, behaving as a single fiber.

To explore this line of reasoning, we calculated the genome-wide average contact frequencies for *cis* as well as *thom* contacts between pairs of loci, i and j (with *i* and *j* being on different homologs for *thom*) as a function of genomic separation (s = |i-j|) measured in base pairs (bps) (Fig. 2Bi top): P_thom_(s) and P_cis_(s) (Supplementary Material). In fact, when the P_thom_(s), is plotted as a function of genomic separation, it is clear that *thom* contacts can occur at all genomic separations. Remarkably, P_thom_(s) peaks at the smallest genomic separation (s = 1 kb) in near perfect registration, with *thom* contacts being as frequent as contacts in *cis* at s = 8 kb. Moreover, the ratio P_thom_(s)/P_cis_(s) is as high as 0.5 to 0.7 for genomic separations of 1 to 10 kb and only gets higher, approaching 1.0, with increasing genomic separation (Fig. 2Bi, bottom). This high ratio indicates that a locus on one chromosome interacts with another locus nearly as often in *thom* as it does in *cis,* especially for s > ~100 kb (Fig. 2Bi, iii, shaded). The simplest interpretation of this observation is that, overall, homologs are aligned in good register genome-wide, almost as a railroad track. We note, however, three caveats. First, our analysis can only capture *thom* interactions that are accessible by Hi-C technology; for example, repetitive regions of the genome cannot be mapped by Hi-C in either *thom* or *cis.* Second, our studies assume that *cis* and *thom* interactions are equally tractable. Third, as our Hi-C studies are a population assay, we cannot rule out cellular heterogeneity in the degree of pairing.

Besides the general distance-dependent decay of *cis* and *thom* contact frequency, our Hi-C maps also revealed a rich structure of *thom* interactions, including well-defined *thom* domains, at genomic separations as small as tens of kilobases, as well as plaid patterns of contacts far off the diagonal, at genomic separations as large as tens of megabases and corresponding to compartments (*2*) (fig. S3A). Consistent with railroad track pairing, we found strong concordance between the *thom*, *cis*-maternal, and *cis*-paternal Hi-C maps in terms of the positions and sizes of domains (Fig. 2C, with Hi-C diagonal positioned horizontally; fig. S3B, C): 81.5% and 89.1% of the domain boundaries in the *cis* and *thom* maps appeared in the *thom* and *cis* maps, respectively (Fig. 2D). Overall, the strong concordance between *thom, cis*-maternal, and *cis*-paternal Hi-C maps indicated a high level of registration between paired homologs.

Looking more closely at our *thom*, *cis*-maternal, and *cis*-paternal Hi-C maps, we discovered that some *thom* domains lacked a prominent signal along the diagonal (Fig. 2C, lower panel showing subtraction Hi-C map; fig. S3D), suggesting an overall looser pairing. This absence of the diagonal contributed to the lower values of P_thom_(s)/P_cis_(s) at genomic separations of s < ~100 kb (Fig. 2Bi, ii unshaded), and clearly demonstrated that pairing was not uniform across the genome. We quantified this variation of pairing via a pairing score (PS), defined as the log_2_ average *thom* contact frequency near the diagonal, that is, where both reads of a read pair lie within a ±12 kb window around a given 4 kb bin, and compared that score to an analogous score for *cis* contacts (CS) (Fig. 2E; Supplementary Material). Thus, as Hi-C data reflect the frequency with which interacting genomic regions colocalize, the PS served as a proxy in our analyses for the relative tightness or looseness of pairing across a chromosomal region. Figure 2E illustrates how dramatically the PS can vary along the chromosome, dipping most noticeably when the *thom* diagonal is missing from the central region of a domain. In line with this and compared to scores for *cis* contacts, PS values for loci within 4 kb (one bin) of domain boundaries are overall much higher than those for loci greater than 28 kb from the nearest boundary, consistent with the lack of diagonals coinciding with the central regions of domains (Fig. 2G) and pointing to tighter pairing at domain boundaries. Lack of a diagonal may reflect any number of structures, including imprecise and/or loose pairing or even the side-by-side alignment of homologous, yet distinguishable, domains (Fig. 2F, schematics below).

To better understand genome-wide variation in pairing, we examined the PS distribution and noted that it could be approximated by two normal distributions. These distributions suggested two classes of loci, one consisting of more tightly paired (higher PS) loci and the other consisting of more loosely paired (lower PS) loci, defined using only a single cut-off (PS = −0.71) (fig S4B). While such a deconvolution likely oversimplifies the reality of pairing, we nevertheless used it to bootstrap our investigation forward. Specifically, we divided the Hi-C amenable portion of the whole genome into regions of tight and loose pairing by first classifying each domain as either tightly or loosely paired based on its PS, and then merging consecutive domains of the same pairing type into one region (fig. S4A, B; Supplementary Material). According to this classification procedure, ~36% of the genome is loosely paired, and ~64% is tightly paired (fig. S4C). From these tight and loose regions, we then selected those spanning distances large enough for us to conduct our studies (Supplementary Material, fig. S5) and calculated P_thom_(s) and P_cis_(s). Tightly and loosely paired regions differed in the decay of *cis* and *thom* contact frequencies. In tightly paired regions, *thom* contacts at the highest registration (smallest genomic separation, s = 1 kb) appeared as frequent as *cis* contacts at s = ~5 kb and, in loose regions, the frequency of such *thom* contacts matched that of *cis* contacts at s = ~30 kb (Fig. 2H, marked on graph). This indicated that, in loose regions, homologs were aligned less precisely. Surprisingly, we found that regions of tight and loose pairing also differed in their internal organization. This was evident from the different shapes of their P_cis_(s) curves – in tight regions, the P_cis_(s) curve had two modes (Fig. 2H, left), a shallow mode at s < ~30kb and a steep mode at s > ~30 kb, while in loosely paired regions, we observed only a shallow mode (Fig. 2H, right). Drawing from other Hi-C studies, where the presence of a shallow mode followed by steep mode is a signature of domains (*38, 39*), we then further interpreted our *cis* data. In particular, the transition of P_cis_(s) at ~10-30 kb for tightly paired regions suggested that they consisted of a series of relatively small domains, within which pairing may reflect primarily the constraints imposed by tight pairing at the boundaries. In contrast, we did not see a similar transition of P_cis_(s) for loosely paired regions, suggesting that each of these regions constituted a single domain. This distinction between tight and loose regions is also evident from visual inspection of the data (Fig 2F).

Having elucidated the structure of paired homologs and its variations, we next addressed the question of whether homolog pairing may bear a genome-wide relationship to genome function. In particular, we conducted three genome-wide analyses, assessing whether pairing correlates with specific epigenetically defined types of chromatin, A- or B-type compartments, and/or gene expression. With respect to chromatin types, we turned to the five defined by Filion *et al*. (*40*) in *Drosophila*, wherein Polycomb group (PcG) repressed chromatin (H3K27me3 enriched) is dubbed blue, inactive chromatin (lacking epigenetic marks) is dubbed black, heterochromatin (HP1 associated and H3K9me3 enriched) is dubbed green, and active chromatin within enhancers/promoters (H3K36me3 depleted) and gene bodies (H3K36me3 enriched) are dubbed red and yellow, respectively. By comparing the coordinates for chromatin types identified in Kc_167_ cells to the PS track, we found that even a localized survey of 1.5 Mb of the genome easily revealed that low PS regions coincide with inactive (black) and repressed (blue) chromatin types, while active chromatin (yellow and red) is present in regions of high PS (Fig. 3A). Interestingly, these trends were confirmed globally, with active regions enriched for high PS, heterochromatin (green) showing a bimodal distribution, and repressed (blue) and inactive (black) chromatin enriched for low PS (Fig. 3B).

**Fig. 3.**
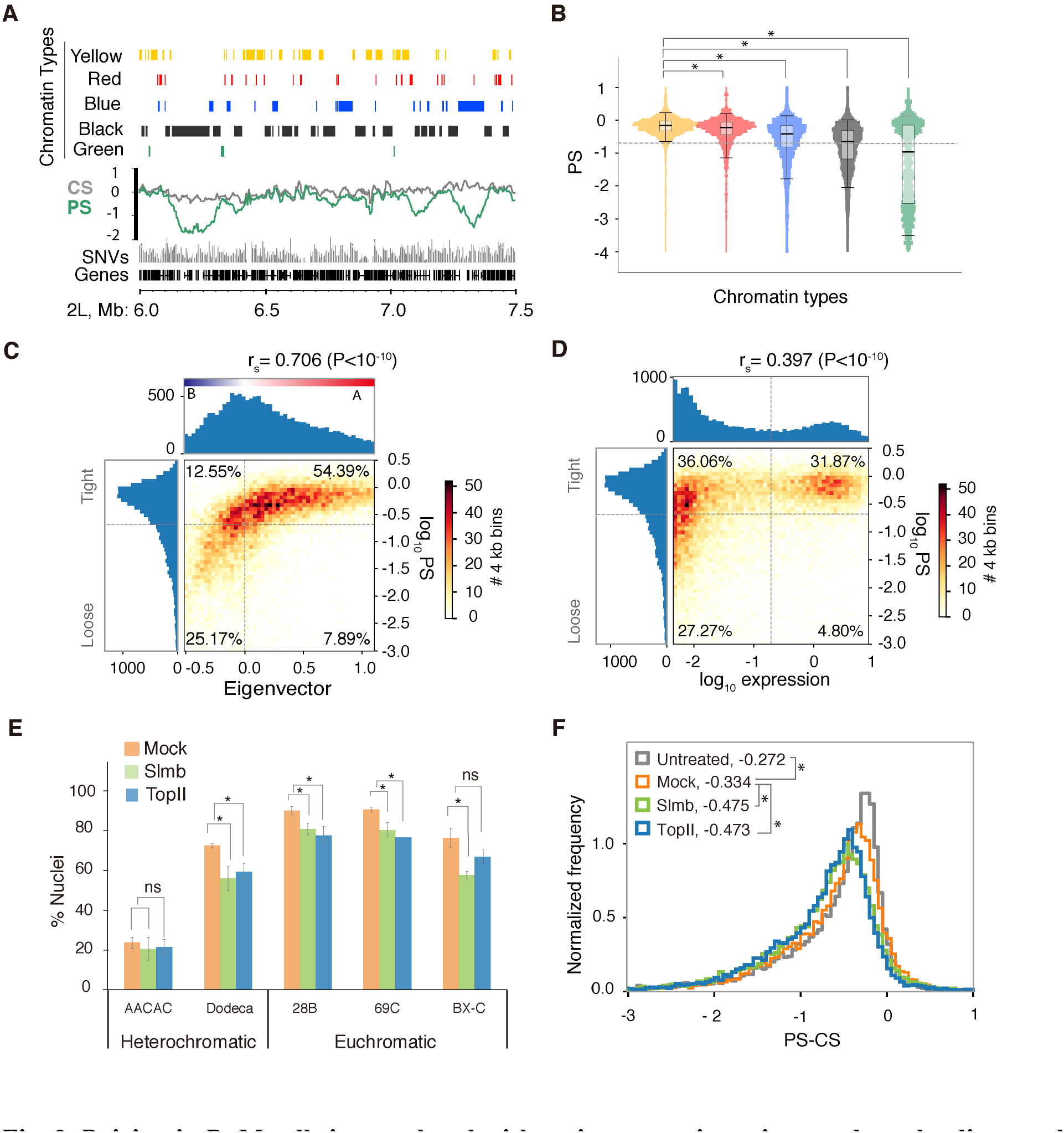
Pairing in PnM cells is correlated with active genomic regions and can be disrupted by RNAi. (A) Pairing scores (PS) and *cis* scores (CS) within a 1.5 Mb region of 2L shown in a genome browser compared to chromatin types identified in Kc_167_ cells (*40*) as well as with SNVs in PnM cells. (B) Normalized distributions of PS within regions of different chromatin types. Dashed line, threshold between tight and loose pairing. *, P<1^-10^, Mood’s test against yellow chromatin. (C) Distribution of PS values relative to the eigenvector shows that pairing is correlated with compartmentalization, A-type compartments being almost exclusively tightly paired. (D) Distribution of PS values relative to gene expression in PnM cells shows that expressed genes are almost exclusively tightly paired. (E) Levels of pairing (quantified and displayed as in Figure 1D) in PnM cells after Slmb and TopII knockdown showed ~10% reduction as compared to the control (mock) at all loci (*, P<=0.05, unpaired t-test) except the AACAC satellite repeat and, in the case of TopII knockdown, at BX-C (ns, non-significant; error bars, s.d for three biological replicates; for N > 100 nuclei/replicate). (F) After Slmb and TopII knockdown, the aggregated pairing score (APS) values were reduced by 0.203% and 0.201%, respectively, as compared to untreated sample (p < 0.001) and 0.141% and 0.139%, respectively, as compared to mock (p < 0.001). The 0.062% reduction in mock as compared to untreated samples was also significant (p <0.001). P-values determined using bootstrapping (Supplementary Material).

Next, we examined the relationship between pairing and the 3D spatial compartmentalization of active and inactive chromatin (*2*). Here, we observed a strong correlation between high PS values and the *cis* eigenvector track (a measure of compartments as determined from Hi-C maps) in individual genomic regions (fig. S7) as well as genome-wide (r_s_ = 0.71, p<10^-10^; Fig. 3C). Regions with high PS values and thus likely to be tightly paired were in predominantly active A-type compartments (54.4% of mappable genome) as versus inactive B-type compartments (12.6%). Conversely, regions with lower PS values and thus likely to be loosely paired were more often in B-type (25.2%) as versus A-type (7.9%) compartments (Fig. 3C). In short, homolog pairing was correlated with compartmentalization of the genome, and active A-type compartments were more likely to be tightly paired. As compartmentalization of the genome into active and inactive compartments may be independent of domain formation by loop extrusion (*38*), these observations may suggest that, compared to domains, pairing may be as, and perhaps even more, related to compartments and the epigenetic states governing them.

Finally, we performed RNA-seq to assess gene expression in PnM cells and found that pairing correlates with gene expression in individual genomic regions (fig. S7) as well as genome-wide (r_s_ =0.40, p<10^-10^; Fig. 3D); regions that are expressed are predominantly tightly paired and have high values of PS (31.9% mappable genome), with only a small percentage (4.8%) of expressed loci being loosely paired (Fig. 3D). On the other hand, lack of expression is not predictive of the degree of pairing; lowly-expressed regions can be associated with either high or low PS values (36.1%, and 27.3%, respectively). These analyses indicated that most active regions (A compartments or regions of high expression) are tightly paired, while repressed and inactive regions demonstrated a variable degree of pairing. In brief, all three approaches argue strongly that pairing bears a genome-wide relationship to genome function.

As many loci in Drosophila have been documented to support transvection, we categorized the commonly recognized autosomal loci with respect to whether they resided in tightly or loosely paired regions in the PnM genome. Fifteen of the seventeen loci for which transvection or related pairing-related phenomena have been reported are associated with tight pairing (with two also associated with loose pairing) and, thus, fall in the upper quadrants of Figure 3D (table S3). Excitingly, this list includes the Antennapedia and Bithorax complexes (ANT-C and BX-C), which include HOX genes critical for body segmentation in *Drosophila* (*41*). While they are known to interact (*7, 42*), our data explicitly reveal their interaction in *thom (*fig. S8) and show their co-localization within the same compartment.

Our final goal was to determine the potential of PnM cells to develop into a robust system for interrogating the mechanism of pairing. We aimed to determine, first, whether PnM cells are responsive to dsRNA, second, whether knockdown of genes involved in pairing (*11, 43-48*) would affect pairing and, third, whether disruptions of pairing as detected by FISH would be detectable via Hi-C. These issues were key. While previous studies had successfully disrupted pairing in *Drosophila* cell lines, under no circumstance had the pairing in diploid cells been disrupted beyond ~50% (*44, 47*) due probably at least to the incomplete nature of RNAi-directed knockdown and perdurance of gene products. It is also possible that pairing, once established, is not easily disrupted (*44, 49*). Finally, disruptions of pairing might affect primarily those forms that are not amenable to detection by Hi-C; as FISH studies often consider two loci to be paired when the center to center distances of the corresponding FISH signals are as far apart 0.5 to 1.0 μm, it is possible that some changes in pairing cannot be captured by Hi-C.

Excitingly, using RNAi to target two genes known to promote pairing, Slmb (component of SCF^Slmb^ complex; (*45, 47, 48*)) and Topoisomerase II ((TopII;(*44*)), we reduced the corresponding mRNA levels by 75.2 ± 2.8% and 82.5 ± 5.0%, respectively (fig. S9A). Importantly, we observed a concomitant 10.7-12.1% reduction of pairing as assayed by FISH (Fig. 3E; table S4). While this reduction was less than previously reported (*44, 47*), it was significant as compared to mock RNAi trials (P< 0.05) (Fig. 3E). We then generated Hi-C maps for the knockdown and mock conditions, each with about 20 million haplotyped mappable reads (table S2), and found a reduction in P_thom_(s)/P_cis_(s) for both Slmb and TopII RNAi samples at all separations as compared to mock and untreated sample (fig S9B; error bars for each sample fall within lines). Note that, while the values of P_thom_(s)/P_cis_(s) for the mock samples veer below those for untreated controls at genomic separations greater than 100 kb, they nevertheless remain above the values for both RNAi-treated samples. Slmb and TopII knockdowns also produced a change in PS. To quantify this change, we computed the aggregated pairing score (APS) as the mode of (PS-CS) distribution, which summarizes the degree of pairing with a single value (Supplementary Material). As shown in Figure 3F, APS dropped after knockdown of Slmb or TopII, compared to mock, and the untreated sample. These observations were consistent across replicates (fig. S9C) and across tight and loose regions (fig. S10; Supplementary Material). In summary, not only were PnM cells amenable to RNAi, but Hi-C could detect global changes in pairing as a result of the knockdown of pairing factors.

In conclusion, we established a hybrid PnM cell line, which allowed us to use haplotype-resolved Hi-C to distinguish *cis* and *thom* interactions, revealing great detail in the structure of homolog pairing for the first time (Fig 4 A, B), in addition to uncovering a genome-wide correlation with gene expression. Furthermore, we observed a minimum of two forms of pairing (Fig. 4B): a tighter, more precise form consisting of contiguous small domains paired at their boundaries and a looser, less precise form often corresponding to single domains that come together at the domain boundaries. The concordance of loose pairing with domains is consistent with both a transgene-based study (*50*) as well as super-resolution images of juxtaposed domains (*18*). We also examined the relationship between pairing and genome function, discovering that active, expressed regions correlated with tighter pairing. While this finding may suggest that gene activity promotes pairing, it is also possible that pairing facilitates the formation of microenvironments favoring transcription; pairing may promote the entangling of R-loops (Fig. 4B) or enrichment of RNA polymerase, transcription factors (*17, 51, 52*), and insulator elements and associated proteins at domain boundaries (*4, 5, 17, 53, 54*). In contrast, transcriptionally inactive regions were either tightly or loosely paired. Thus, tight pairing may facilitate either activation or gene repression, consistent with studies of transvection (reviewed by (*8-12*)). Note that our findings differ from predictions of a study (*23*) that, in the absence of haplotype-resolved data, was nevertheless able to simulate pairing via the computational integration of Hi-C and lamina-DamID data representing embryos (*7*) and Kc_167_ cells (*55*), respectively. Contrary to our findings, the simulations predicted correlations between active regions and loose or occasionally tight pairing, and between inactive regions and tight pairing. One possible explanation is that these two studies suggest an as yet unexplored mechanism of pairing.

**Fig. 4.**
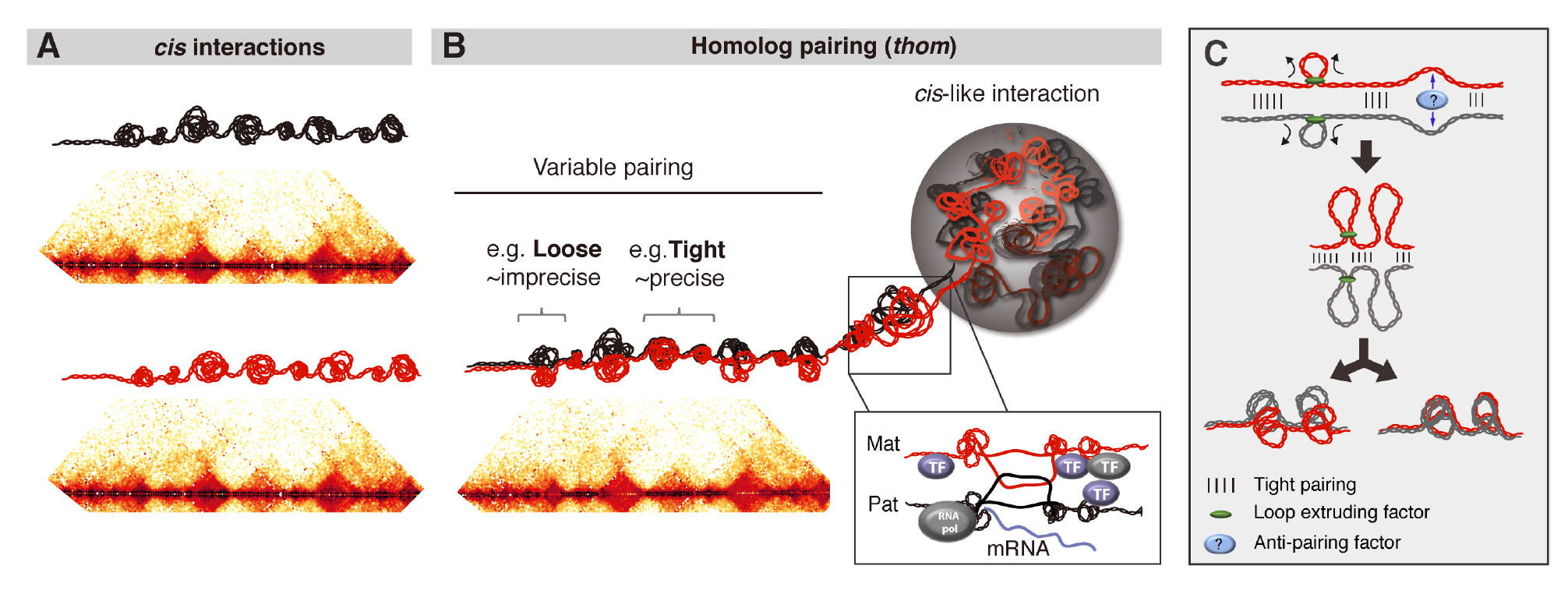
Haplotype-resolved Hi-C in PnM cells distinguished different forms of *thom* and *cis* interactions. (A) *Cis* contact maps for two homologs and schematics depicting possible *cis* interactions. (B) *Thom* contact map demonstrates variable structure of pairing, including tight, precise pairing interspersed with looser, less precise pairing. *Thom* interactions encompass an organization concordant to *cis* behavior, which could facilitate the formation of active transcriptional environments between homologs. (C) Homologous loops may form by extrusion (black arrows) or anti-pairing (blue arrows) between tightly paired regions, which could behave as extrusion barriers. Here, loops could result in *thom* domains that are either railroad-track paired throughout or loosely paired, and *cis*-maternal and *cis*-paternal domain boundaries are expected to be concordant. Note, loop extrusion in mammalian systems is proposed to involve a cohesin ring through which a single chromosome passes (*39*), suggesting that extrusion in the context of pairing may involve the passage two homologs simultaneously. Our model proposes a nonexclusive alternative, in which extrusion of single chromosomes nevertheless achieves *thom* domains.

Loosely paired regions are equally interesting, as they lack a *thom* diagonal. This feature, which was also observed in embryos (Erceg, AlHaj Abed, Golobordko *et al. bioRxiv (37)*), indicates a lack of railroad track pairing (Fig. 2F, schematics below). Importantly, the boundaries of these loosely paired regions are tightly paired, recalling a model that integrates pairing and loop formation (*47*). With respect to loosely paired regions, homologs could be extruded via any number of mechanisms (*39*) or anti-paired between tightly paired regions but still interact by virtue of remaining tightly paired at their loop bases. In this scenario, tightly paired regions could behave as extrusion barriers and become boundaries (Fig. 4C). Lack of a diagonal has also been observed for polytenized chromosomes (*56, 57*), where it may reflect an outnumbering of *cis* contacts by an abundance of *trans* contacts. These observations raise the possibility that under some circumstances there may be a competitive relationship between short-range *cis* and *thom* contacts (*11, 58*) and/or between short-range and long-range *thom* contacts. In this vein, we turn to our observation that loci interacted with a second locus in *thom* nearly as often as they did in *cis*. Indeed, researchers have long speculated about the consequences of providing regulatory regions with a *cis*-*trans* choice (*9, 11*). For example, pairing may enhance co-regulation of allelic regions by enabling transcriptional states to be transferred from one chromosome to another (*14, 59, 60*), while the unpaired state may facilitate allele-specific expression, especially in mammals where homologs pair transiently and in a locus-specific manner ((*61*); reviewed by (*9, 11*)). Homolog pairing may even accomplish what compartments do in both *cis* and *trans*, and what domains do in *cis* (*58*), co-localizing genomic regions to achieve an “economy of control (*60*)”.

## Acknowledgments

We thank the Wu and Mirny laboratories, participants of the Annual Northeast Regional Chromosome Pairing Conferences, the Lieberman Aiden laboratory, F. Bantignies, B. Beliveau, G. Filion, M. Francesconi, E. Joyce, B. Lehner, and J. Rowley for discussions, the TUCF Genomics Sequencing Core Facility for sequencing services, and the Drosophila Genomics Resource Center (NIH 2P40OD010949) for cell lines.

## Author contributions

J.A.A., J.E., and C.-t.W. designed the experiments. A.G. designed Hi-C computational analyses with input from J.E., J.A.A., L.A.M. and C.-t.W. J.D. and B.R.L. provided input in experimental design and data analysis. J.A.A., J.E., A.G., B.R.L., G.F., R.B.M, and W.S., carried out analyses of the Hi-C data. Experimental data were generated by J.A.A. and J.E. J.E. and R.B.M. selected the parental lines. J.A.A. validated and characterized the cell lines, which were made by S.C.N. J.A.A. performed RNAi analyses. J.D., L.A.M., G.F., and B.R.L provided advice on Hi-C analyses. J.A.A. ., J.E., A.G., G.F, L.A.M., and C.-t.W. interpreted the data. J.A.A., J.E., A.G., L.A.M., and C.-t.W. wrote the paper with input from the other authors.

## Data and materials availability

Raw sequencing data and extracted Hi-C contacts has been deposited in the Gene Expression Omnibus (GEO) repository under accession number GSE121256. Hi-C data obtained in this study is available for browsing using the HiGlass web browser (*62*).

## Materials and Methods

### Fly Stocks and Crosses

We selected two highly divergent parental fly lines from the *Drosophila* Genetic Reference Panel (DGRP) (*1*), and set up a cross between (DGRP-057 females x DGRP-439 males (Erceg, AlHaj Abed, Golobordko *et al. bioRxiv* (*2*)). Primary cultures were established from an overnight embryo collection aged for 2 hours (2-14 hr AEL).

### Establishing Primary Culture PnM and clonal cell line

Primary cell line was generated as described previously (*3*). Hybrid ~ 2-14 hour old embryos were collected overnight at 25°C on agar juice plates covered with killed yeast paste. After the embryos were rinsed from the plates and collected in a 50 ml sieve basket they were washed using TXN wash buffer (0.7% NaCl, 0.02% Triton X-100). The TXN was replaced with 50% bleach in water for 5 minutes to dechorionate and surface sterilize the embryos. The embryos were washed extensively with TXN, sprayed with 70% ethanol, and transferred to a sterile tissue culture hood, where they were rinsed once in water and placed in 3 ml medium for 5 minutes (unsupplemented M3 Insect Media (Sigma)). Around 100 μl embryos or more were transferred to a homogenizer (Wheaton 5 ml) and homogenized in 5 ml M3 Insect medium with a loose pestle ¾ of the way until embryos were homogenized, then 5X with a loose pestle all the way, followed by 5X with a tight pestle. The homogenate was transferred to a 15 ml conical tube, and cell debris was pelleted at 500 rpm. The supernatant was transferred to a new tube and cells were pelleted by centrifugation at 1200 rpm, resuspended in fresh media and counted. 8 million cells were plated in 25 cm^2^ T-flasks and grown at 22°C with 5 ml of supplemented M3 media (M3 Insect Media, 10% Fetal Bovine Serum (JRH), 2.5% Fly Extract (DGRC), 0.5 mg/ml Insulin (Sigma), 1:100 Pen-Strep (Gibco)). After a couple of weeks, the media was replaced with conditioned supplemented M3 media (0.2 μm filter-sterilized used media), and was then changed every couple of weeks for a few months. With continued passaging the cells became stable, and immortalized spontaneously. Then, as they divided more regularly, they were trypsinized with TrypLE 1x (ThermoSci) and split at 1:5 into supplemented M3 media every 2-3 days. Establishing a clonal cell line was done using a serial dilution single cell cloning protocol http://www.level.com.tw/html/ezcatfiles/vipweb20/img/img/34963/3-2Single_cell_cloning_protocol.pdf . One of the clonal lines isolated, Pat and Mat line #A1 (PnMA1), is used in this study and is referred to as PnM.

### Cell culture

Kc_167,_ and clone8 (cl.8+) cells were obtained from the Drosophila Genome Resource Center and cultured according to standard protocols at 22°C (see www.flyrnai.org for more details). Hybrid PnM cells were cultured as cl.8+ cells with supplemented M3 media (supplemented M3 media (M3 Insect Media, 10% Fetal Bovine Serum (JRH), 2.5% Fly Extract (DGRC), 0.5 mg/ml Insulin (Sigma), 1:100 Pen-Strep (Gibco)), according to standard protocols at 25°C in small culture dishes (100x 20 mm), with the following modifications: the cells were trypsinized and split at 1:5, or 1:10 every other day, and the media was replaced every day after they are washed with 1X PBS. To ensure the cells were in log phase for any experiment, they were split 12-18 hrs earlier.

### Genomic DNA extraction, and PnM genome sequencing

Around 1-2 million PnM cells were used to isolate Genomic DNA using Qiagen DNeasy Blood & Tissue Kit, and ~ 0.5μg genomic DNA was used for the generation of an Illumina TruSeq Nano DNA Library. The library was 150 bp paired-end sequenced using Illumina HiSeq2500 at TUCF Genomics core.

### FACS

Cells were split a day prior and then 1-3 million cells were harvested and fixed with 95% ethanol, washed with 1X PBS and stained using FxCycle PI/RNase staining solution (Life Technologies) for 30 min at room temperature. Cell populations were assayed based on DNA content to determine their cell cycle profile using an LSR II Analyzer at the HMS Immunology flow cytometry core.

### Preparing metaphase spreads and karyotyping

Metaphase cells were prepared using protocols adapted from published methods (*4*). 150 μl of colchicine was added to 5 ml cells in culture at a concentration of 30 μM for 45 min prior to fixation and spread preparation. Cells were spun down at 1200 rpm, washed with 1X PBS and resuspended in 10 ml 1% sodium citrate slowly, while vortexing regularly. The cells were incubated at room temperature for 30 minutes. Then 1 ml of cold fixative was added (3:1 methanol: glacial acetic acid solution) while vortexing gently, cells were spun down, and washed three more times in 10 ml of the same fixative. Finally, cells were resuspended in 1 ml of the fixative and dropped at a height of 5 inches or more onto a glass slide under humidified conditions. The slide was allowed to dry in a humidified chamber and then washed in 70%, 90% and 100% ethanol successively. For long-term used they were stored at 4°C, in 1X PBS. In order to examine the karyotype, slides were mounted with Slowfade Gold Antifade with DAPI (Invitrogen). The spreads were then examined with a Nikon Eclipse Ti at 60X. Karyotype was examined for PnM cells for N=50 from 2 different metaphase spread preparations, and showed the cell line to be male and diploid, apart from chromosome four. The fourth chromosome was 7% diploid, since it was either monosomic (~71%) or trisomic (~22%). Given its ploidy, and that it is largely heterochromatic, with an average 1 SNV per kb (compared 5 SNVs per kb for the second or third, table. S1), it was not included in the haplotype-mapping and analysis.

### FISH probes

Heterochromatic repeat regions were assayed using previously described FISH probe sequences (*5, 6*), and used to assay FISH loci localization as shown previously (*7, 8*). The probes used were synthesized by Integrated DNA Technologies (IDT) as follows: Alexa647-359-X: /5Alex647N/-GGGATCGTTAGCACTGGTAATTAGCTGC, Atto565-AACAC-II: /5Atto565N/ AACACAACACAACACAACACAACACAACACAACAC, and Alexa488-dodeca-III: /5Alex488N/-ACGGGACCAGTACGG. The euchromatic Oligopaints FISH probes used in this study, 16E, 69C and 28B as well as HOPs probes were described previously (*8-10*) and were generated using the T7 method (*11*). The oligopaint libraries used as template are described in table. S5 and are amplified using the primers shown in table. S6. Secondary probe binding sites were added to oligopaints mainstreets, and T7 sequences to backstreets by touch-up PCR. For detecting the euchromatic oligopaints, a secondary fluor-tagged probe sequence complementary to the mainstreet, was co-hybridized with the primary probe. Homolog specific Oligopaints (HOPs) probes include at least one SNV location per oligpaint, and are used to distinguish either 2L or 3L homologs in the hybrid line by targetting a 2 Mb sub-telomeric region on either Chr2, and Chr3. The secondary, dual labeled probes used for detection of euchromatic targets are ordered from IDT and are as follows: Secondary1: /5Alex488N/CACACGCTCTTCCGTTCTATGCGACGTCGGTGagatgttt/3AlexF488N/, secondary5: /5Atto565N/ACACCCTTGCACGTCGTGGACCTCCTGCGCTA/3Atto565N/, and secondary6: /5Alex647N/TGATCGACCACGGCCAAGACGGAGAGCGTGTGagatgttt/3AlexF647N/.

### Metaphase FISH

Metaphase FISH was done as described previously (*4*). Slides from the metaphase spread preparation were rehydrated in 1X PBS for 5 minutes, and denatured in 67% formamide/2X SSCT at 80°C for 90 seconds, followed by washes in ice-cold 70%, 90% and 100% ethanol. 20 pmol of HOPs were co-hybridized with 40 pmol of secondaries without any additional denaturation at 42°C overnight. The remainder of the protocol is the same as in the interphase FISH protocol.

### Interphase FISH

Fluorescence *in situ* hybridization was done as in described previously (*11*). A cell suspension at 0.5-1 million cells per ml was allowed to settle on poly-l-lysine coated slides for a few hours, washed with 1X PBS, then fixed in 4% paraforlamdehyde and washed in 1X PBS again. The slides were either used for FISH immediately or stored in 1X PBS at 4°C. Just before FISH, the cells were permeabilized by incubating in 0.5% PBST for 15 minutes, and 10 minutes in 0.1 M HCl. The slides are then washed in 2X SSCT for 5 min (0.3M sodium chloride, 0.03M sodium citrate, 0.1% Tween-20), and 50% formamide/2X SSCT. FISH slides were incubated in 50% formamide/2X SSCT 60°C for 20 minutes. FISH probes were added in a hybridization solution of 10% dextran sulphate/2X SSCT/50% formamide and 100 pmol of heterochromatic, or 50 pmol euchromatic probes, 16E, 28B, 69C and BX-C per hybridization. The slides were then denatured by placing them on a heat block at 80°C for 3 minutes and allowed to co-hybridize overnight at 42°C with 40 pmol of secondary probe. Following hybridization, slides were washed in 2X SSCT at 60°C for 20 minutes, 2X SSCT at room temperature for 5 minutes, and 0.2X SSC at room temperature for 10 minutes before being mounted using Slowfade Gold Antifade with DAPI (Invitrogen) and imaged. To quantify level of pairing for the X, second and third chromosomes at the same time, probes were co-hybridized for 16E-X, 28B-II, 69C-III FISH probes or 359-X, AACAC-II, and dodeca-III. To score cells that are non-replicating, and since the cell line is male, we use probes targeting the X chromosomes, either euchromatic target 16E or heterochromatic target, 359, and consider the pairing only in cells with one X-specific FISH signal per nuclei.

### Image acquisition and analysis

All images were obtained using Nikon Eclipse Ti microscope with a 60X oil objective and Nikon ND acquisition software. The raw TIFF files obtained were analyzed using custom-written MATLAB scripts (*7*) and later adapted (*4*) for measuring different properties such as the number of FISH dots per nucleus. All uniquely identifiable foci of fluorescent signal (above background) were counted as FISH signals. The number of FISH foci were also counted manually to confirm consistency and determine the degree of localization of FISH foci in 3D by eye. Homologous loci were considered paired if FISH signals targeting the loci co-localized (i.e., gave a single signal) or exhibited a center-to-center distance of ≤ 0.8 μm.

### Immunostaining to determine cell type and Mitotic index

PnM cells were fixed with 4% formaldehyde for 20 minutes, following previously published protocols (*4*). In order to determine the mitotic index for the PnM hybrid clone, a primary antibody against phosphohistone H3 (P-H3; rabbit used at 1:100; Epitomics) was used for immunofluorescence in a 1X PBS buffer. A Cy3-conjugated anti-rabbit secondary antibody (Jackson ImmunoResearch Laboratories) was used at 1:100. Mitotic index for PnM cells was determined to be 2.48% ±0.78, N=1300.

In order to determine cell type as previously described (*12*), PnM were immunostained using rabbit anti-dMef2 antibody, which was a gift from Bruce Paterson (*13*). The antibody was used at 1:1000, with a Cy3-conjugated anti-rabbit secondary antibody at 1:100 (Jackson immunoresearch). The cells expressed dMef2 exclusively, and are most likely of mesodermal origin, as they tested negative for other cell type markers including D-E-Cadherin ((Rat)-1:5 (Hybridoma Bank, Iowa), epithelial cell marker, Nile red solution (Sigma; 1% stock in DMSO diluted to 1:5000), fat cell marker, and HRP-(Jackson immunoresearch (Rhodamine conjugated) 1:200), a nerve cell marker.

### dsRNA synthesis and RNAi Treatment

Synthesis of dsRNA was carried out according to standard protocols (see www.flyrnai.org for more details). Control cells were treated with blank dionized water. Primers used for dsRNA synthesis are listed in table. S7. The dsRNA was administered to the cells using a calcium phosphate transfection kit (Invitrogen) in 60-mm well plates. Kc_167_ cells were seeded at 2 million cells /ml and treated with 15 μg dsRNA and harvested after 3 days. PnM cells were seeded at 1 million cells /ml and treated with 30 μg dsRNA on the first and third day, and then harvested on the fourth. Cells from the knockdowns were counted and an aliquot was taken for RNA isolation, and once knockdown of mRNA was confirmed cells were split to be fixed for Hi-C, and for FISH. Two biological replicates were processed for each treatment. Knockdowns in Kc_167_ cells were used as a control for the knockdown experiment, but were not processed for Hi-C experiments.

### qPCR

Quantitative PCR was used to assay efficiency of RNAi knockdowns according to standard techniques. Total RNA was isolated from cells using a Qiagen RNeasy Plus kit and then converted to cDNA using SuperScript VILO cDNA synthesis kit (Invitrogen) for RT-PCR. Primers for qPCR were designed using Primer3 website (http://bioinfo.ut.ee/primer3-0.4.0/) and are listed in table. S8. Reactions were set up according to recommended protocol using iQ SYBR Green Supermix (BioRad) and run on BioRad CFX Connect Real-Time System at an annealing temperature of 58°C. BioRad software determined CT values for qPCR reactions, and the level of knockdown is determined using the 2^(*-∆∆CT*)^ method (*14*). The level of knockdown was normalized to two controls; Act5c, and RP49.

### *In situ* Hi-C protocol

This protocol was adapted from a previously published protocol (*15*) with modifications. Unless otherwise specified, 75T flasks of PnM cells were cultured to 70% confluency, then washed with FBS-free Schneider’s medium, and then crosslinked with 1% formaldehyde for 10 min at room temperature. Fixation was quenched with 1 M glycine solution, and the cells were scraped gently off the flask, spun down at 1200 rpm, and washed once more with 1X PBS. Supernatant was removed, and cells were resuspended in 1X PBS and then ~2.5 million cells were counted to be used for the protocol. Nuclei were permeabilized with ice cold lysis buffer (10 mM Tris-HCl pH 8.0, 10 mM NaCl, 0. 2% Igepal supplemented with 5X Complete, EDTA-free Protease inhibitors (Roche). DNA was digested with 500 units of DpnII, and the ends of restriction fragments were labeled using biotinylated nucleotides and ligated in a small volume. After reversal of crosslinks, ligated DNA was purified and sheared to a length of ~700 bp with QSonica sonicator (30% power, 30 sec on, 30 off for 20 minutes, at which point ligation junctions were pulled down with Dynabeads MyOne Streptavidin beads (Invitrogen) and prepped for Illumina sequencing. Aliquots at different steps were taken to measure the concentration with Qubit dsDNA HS assay kit, and run on a 2% agarose gel with SYBR Gold nucleic acid gel stain (1:10000) to ensure quality control of the sample prepared, and to ensure efficient digest, ligation, and binding to beads. For library preparation, we used BioNEXTFLEX barcode-6 (Bioo Scientific) and followed manufacturer recommendations, and amplified our final library with Q5 HIFI Hot Start High Fidelity PCR and Biooscientific primer mix for 6 cycles on the beads. The final product was then diluted to 250 μl with 1 mM Tris-HCl, and separated on a magnet then transferred the supernatant to a new tube and purified with 0.7X AMPure XP bead (Beckman Coulter).

Incubation times are extended to 15 minutes to maximize library recovery. To remove traces of short products, we resuspend beads in 100 μl of 1X Tris buffer and add another 70 μl of AMPure XP beads. Mix by pipetting 20x and incubated at room temperature for 15 min, and separated for another 15 min, and continue with the washes and drying as previously described. Once the beads are dry, we resuspended in 20 uL 1X Tris-HCl, incubated for 15 minutes, then separated for another 15 min on the magnet and transferred the final library to a fresh tube. Two replicates were prepared per sample. The library quality was assessed using the High Sensitivity DNA assay on a 2100 Bioanalyzer system (Agilent Technologies). Then were 150 bp paired-end sequenced using Illumina HiSeq2500 at TUCF Genomics core. PnM untreated sample was sequenced in 4 lanes, while each RNAi treatment was sequenced in one lane.

### RNAseq

Total PnM RNA was purified from ~8 million cells per replicate using TRIzol (Life Technologies), followed by chloroform extraction, DNase treatment with RNase free DNase I recombinant (Roche), and clean up with RNeasy Mini kit (Qiagen). The quality of total RNA was determined using Agilent RNA 6000 Pico assay on a 2100 Bioanalyzer system (Agilent Technologies). RNA-Seq libraries were prepared using NEBNext Poly(A) mRNA Magnetic Isolation Module and NEBNext Ultra Directional RNA Library Prep Kit for Illumina according to the manufacturer’s instructions. Poly(A)+ RNA was enriched from 1 μg of total RNA, fragmented for 10 minutes at 94°C, and reverse transcribed in the first strand cDNA synthesis with random primers. After adaptor ligation, cDNA was size selected between 400-600 bp with Agencourt AMPure XP beads, and amplified for 12 PCR cycles. The library quality was assessed using the High Sensitivity DNA assay on a 2100 Bioanalyzer system (Agilent Technologies). RNA-Seq libraries corresponding to three biological replicates were 150 bp paired-end sequenced with Illumina HiSeq2500 at TUCF Genomics core.

## Bioinformatic analysis

### The construction of diploid PnM fly genome

We generated the diploid genome (hybrid PnM genome) using (i) the homozygous autosomal PnM SNVs, (ii) the heterozygous phased autosomal PnM SNVs and (iii) the homozygous chrX PnM SNVs.

After sequencing the F1 PnM cell line at the average coverage of 396 reads per base pair [https://www.ncbi.nlm.nih.gov/pubmed/29096012], we used *bcftools* to detect the sequence variation of this library. We obtained high-quality normalized sequence variants using the following:

1. ‘seqtk trimfq’ to trim low-quality sequences, BWA mem, to align whole genome paired-end reads against the reference dm3 genome, and ‘samtools --rmdup’ to remove aligned PCR duplicates
2. ‘bcftools pileup --min-MQ 20 --min-BQ 20’ to pile alignments along the reference genome.
3. ‘bcftools call’ to call raw sequence variants from the pileups.
4. ‘bcftools norm’ to normalize raw sequence variants.
5. ‘bcftools filter INFO/DP > 80 & QUAL > 200 & (TYPE="SNV" | IDV > 1)’ to select only high-coverage high-quality normalized sequence variants using.

The two inbred parental fly lines (057 and 439) were sequenced at the average coverage of 118, and 117 reads per base pair, respectively, and treated similarly to detect sequence variation in their libraries as described in Erceg, AlHaj Abed, Golobordko *et al. bioRxiv* (*2*).. Using the variants from PnM, and the two paternal lines, we then phased heterozygous PnM variants using ‘bcftools isec’. Then, we picked high-confidence variants on the maternal autosomes by selecting heterozygous PnM variants that were present among maternal 057 variants and absent among paternal 439 sequence variants (both homo- and heterozygous); the high-confidence paternal variants phasing was selected in an opposite manner. Since PnM is a male line, for chrX, we considered only homozygous, high-quality variants detected in the PnM cell line. To reconstruct the consensus sequence of the paternal copy of chrX, we kept only homozygous variants detected in the paternal 439 fly line. Finally, we reconstructed the sequence of the PnM cell line with ‘samtools consensus’, using (a) the homozygous autosomal PnM SNVs, (b) the heterozygous phased autosomal PnM SNVs and (iii) the homozygous chrX PnM SNVs. Overall, WGS confirmed PnM cell line hybrid status, apart from uncovering a 24.6 Mb partial uniparental disomy of the right arm of chromosome 3 (chr3R). We adjusted our downstream analyses to take that into consideration.

## Hi-C data analysis

### Mapping and parsing

Using the standard mode of *seqtk trimfq* v.1.2-r94 [https://github.com/lh3/seqtk], we trimmed low-quality base pairs at both ends of each side of sequenced of Hi-C molecules. This was followed by mapping the trimmed sequences to either, the reference dm3 genome, or the newly constructed dm3-based diploid genome (hybrid PnM genome) using *bwa mem* v.0.7.15 [https://arxiv.org/abs/1303.3997] with flags -SP.

We then used the *pairtools parse* command line tool (https://github.com/mirnylab/pairtools), to extract the coordinates of Hi-C contacts, kept read pairs that mapped uniquely to one of the two homologs, and used the standard mode of the *pairtools dedup* command line tool to remove PCR duplicates.

Detailed methods on haplotype-Hi-C mapping, and estimating the percent of homolog misassignment was discussed in Erceg, AlHaj Abed, Golobordko *et al. bioRxiv (2).*

### Contact probability P(s) curves

To calculate the functions of contact frequency P(s) against genomic separation (s), we used unique Hi-C pairs, and grouped genomic distances between (10bp −10Mb) into ranges of exponentially increasing widths, with 8 ranges per order of magnitude. We found the number of observed *cis*- or *trans*-homolog interactions, within this range of separations, and divided it by the total number of all loci pairs separated by such distances.

### Pairing score

We introduced a genome-wide track called pairing score (PS), to measure the strength of the diagonal and to characterize the degree of pairing between homologous loci across the whole genome. The PS of a genomic bin is log_2_ of average *trans*-homolog IC contact frequency between all pairs of bins within a window of +-W bins. For each genomic bin i, its PS with window size W is defined as:

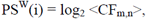

averaged over bins *m* and *n* between i-W-th and i+W-th genomic bins on different homologs of the same chromosome.

As a control measure, we complemented the PS with a *cis* score (CS), which is calculated the same way as the PS, but over the *cis* contact map. Comparing the PS and CS value in a given bin reveals if the local variability in the PS is due to a change in homologous pairing in 3D (which affects the PS, but does not affect the CS) or due to a theoretically possible local deviation from the equal visibility assumption of IC (which would affect both PS and CS equally). Using this definition, the PS quantifies only contacts between homologous loci and their close neighbors and not non-homologous loci on homologous chromosomes.

We chose the window size W to be a balance between specificity and sensitivity. Increasing the window size increases sensitivity, accumulating contacts across more loci pairs, while smaller windows favors specificity, allowing to see smaller-scale variation of homologous pairing. For our contact maps with a 4kb resolution, using a 7×7 bin window (W=3) provided a balance between specificity and sensitivity.

We interpreted the obtained PS tracks using a simple assumption: if a genomic locus was paired with its counterpart on the homologous chromosome in 100% of cells, the frequency of the trans-homolog contacts (i.e. PS) around the locus should be equal to the frequency of corresponding short-distance *cis* contacts (i.e. CS).

### Insulation scores

Using the package *cooltools diamond_insulation* (https://github.com/mirnylab/cooltools) we calculated the tracks of contact insulation score. The method used is based on the algorithm described in (https://doi.org/10.1038/nature14450) and adjusted in (https://doi.org/10.1016/j.cell.2017.05.004).

We calculated the insulation score as the total number of normalized and filtered contacts formed across that bin by pairs of bins located on the either side, up to 5 bins away, for each bin in our contact map binned at 4 kb resolution. Then we normalized the score by its genome-wide median. To find insulating boundaries, we detected all local minima and maxima in the log_2_-transformed and then distinguish them by their prominence (Billauer E. peakdet: Peak detection using MATLAB, http://billauer.co.il/peakdet.html). The insulating boundaries were the detected minima in the insulation score, corresponding to a local depletion of contacts across the genomic bin. We empirically found that the distribution of log-prominence of boundaries has a bimodal shape. Based on that, we selected all boundaries in the high-prominence mode above a prominence cutoff of 0.1 for Hi-C mapped to the reference dm3 map, and a cutoff of 0.3 for allele-resolved Hi-C maps. We called the insulating boundaries in *trans*-homolog contact maps using the same approach, and requiring a minimal prominence of 0.3. Finally, we removed boundaries that are adjacent to the genomic bins that were masked out during IC.

We evaluated the similarity of insulating boundaries that were detected in the *cis* and *trans*-homolog contact maps by calculating the number of overlapping boundaries, and allowing for a mismatch of up to 4 genomic 4 kb bins (16 kb total) between overlapping boundaries to account for the drift caused by the stochasticity of contact maps.

### Detecting tightly and loosely paired genomic bins

We used the genome-wide PS track to classify each genomic bin as either tightly or loosely paired. We noticed that the genome-wide distribution of PS (fig. S4B) showed a well-pronounced peak at higher values of PS and a tail extending into the lower values of PS. We interpreted this distribution with a model, where each bin can be either tightly or loosely paired with a homologous locus on the second chromosome. In this model, tightly paired loci showed high PS values, producing the peak on the genome-wide distribution of PS, while loosely paired loci had lower PS values, giving rise to the tail of the PS distribution. We separated the peak from the tail of the distribution by fitting it with a sum of two Gaussians (fig. S4B); to stabilize the fitting procedure, we also clipped PS values below −3. The probability densities of the two Gaussians become the same at PS = −0.71. Thus, we classified all genomic bins with PS < −0.71 as loosely paired (since they are more likely to belong to the low-PS Gaussian) and the bins with PS > −0.71 as tightly paired.

### Dividing the genome into regions of tight and loose pairing, and P(s)^tight^ and P(s)^loose^ curves

A close examination of the PS track revealed two important features: (i) the fly genome is divided into regions that demonstrate consistently high, relatively similar, values of PS, followed by extended regions where PS dips into lower values, (ii) switching between high- and low-PS regions seemed to occur often around insulating boundaries.

We used the PS and insulating boundaries to determine the precision of pairing in the genome, examine the variation of pairing more closely, and the internal organization of tight and loose pairing. First, we divided the genome into regions between pairs of consecutive boundaries (as detected in reference-mapped, i.e. not allele-resolved Hi-C data). Second, we classified each of these regions as either tight or loosely paired, depending on the number loosely paired bins in that region. Because the dips in the PS track tend to be gradual, with PS being noticeably low only further away from the boundaries, we considered the cutoff of 25% of loosely paired bins per region to be sufficient to call the whole region as loosely paired. Finally, we noticed that this method occasionally split single loosely paired regions into a few smaller ones, presumably due to false positive boundary calls. To mitigate this issue, we merged consecutive loosely and tightly paired regions into larger ones if the boundary bin between them was classified the same way. We then used the detected regions of loose and tight pairing to calculate scaling curves P_cis_(s)^loose^, P_cis_(s)^tight^, P_thom_(s)^loose^ and P_thom_(s)^tight^. We calculated these curves using the same method as for the genome-wide P_cis_(s) and P_thom_(s), but only considering pairs of loci within the same region.

For each type of pairing, we calculated P(s) curves over large regions with a relatively narrow size range of 200-400kb, or 100-200kb (fig. S5). In tightly paired regions (fig. S5A), the P_cis_(s) and P_thom_(s) curves showed two modes: (i) a shallow mode at shorter separations <30kb, where P_thom_(s) was noticeably lower than P_cis_(s) (at 1-3kb, P_thom_(s)/P_cis_(s)) is 0.66, and where both decayed relatively slowly with distance (P(s) ~ s^α^, α~0.25-0.5); and (ii) a steep mode at larger separations >30kb, where the two curves were almost equal (P_thom_(s)/P_cis_(s)= 0.93) and decayed more rapidly (P(s) ~ s^α^, α~1.0-1.5) (Fig. 2H, left panel). In loosely paired regions, P_thom_(s) and P_cis_(s) had only one shallow mode, where P_thom_(s) < P_cis_(s) at all separations, (Fig.2H, right panel); and both curves decayed slowly (P(s) ~ s^α^, α~0.25-0.5). We observed similar behavior of P_cis_(s) and P_thom_(s) in regions of 100-200kb (fig. S5B).

In *cis*, the existence of the shallow mode in the P_c_(s) curves reflects formation of domains, as the slow decay of CF with distance (100x increase in distance leads to only 10x drop in CF) is inconsistent with random folding of the chromatin (*16, 17*). Importantly, the transition into the steep mode occurs at the average domain size (*16*). Thus, our results suggested that an individual tightly paired region in *cis* consists of a series of smaller domains (~10-30Kb), while an individual loosely paired region could correspond to a single domain.

We used the deviation of P_thom_(s) from P_cis_(s) in tightly and loosely paired regions to infer the precision of homologous pairing. If two homologous chromosomes were connected with each other at every base pair (either by direct base-pair-complementarity or by a sequence-specific binding protein or other mechanism), P_thom_(s) would equal P_cis_(s) at all separations. However, if homologous loci were linked with each other intermittently, pairs of loci at shorter distances would contact less frequently in *thom* than in *cis* (P_thom_(s)< P_cis_(s)), and at sufficiently large separations pairs of loci would contact each other as often in *thom* as in *cis* (P_thom_(s) = P_cis_(s)). In our data, in tight regions, *thom* and *cis* contacts approached each other in frequency at s = ~10-30kb, and, in loose regions, only for loci located on the ends of the region (i.e. at s ~150 kb for 100-200 kb regions (fig. S5) and at s~300 kb for 200-400 kb regions). Thus, we concluded that our data was consistent with a model, where tightly paired regions are connected in *thom* every ~10-30 kb, within the average domain size in these regions, and probably at domain boundaries, and loosely paired regions are connected only at their boundaries. This allowed us to hypothesize that (a) the difference between tightly and loosely paired region is due to higher frequency of *thom* connections, in tightly paired regions (b) pairing at loose regions is affected by pairing at the flanking tight regions. To conclude, within our resolution limit (~16 kb), and given that in tightly paired regions, *thom* contacts at the highest registration appeared as frequent as *cis* contacts at s = ~5 kb and, in loose regions, the frequency of such *thom* contacts matched that of *cis* contacts at s = ~30 kb (Fig. 2H), pairing in tight regions is more precise, likely due to more frequent connections between homologs within tight regions, possibly at domain boundaries. Furthermore, we found that *thom* contact frequency approached 93.2% of *cis* in tightly paired regions (at s=100-300kb) (defined as a geometric mean of thom/cis CF at s=100-300kb), suggesting that as many as 93.2% of cells in our sample exhibiting pairing at those regions (fig. S5). The fraction of cells without pairing, could represent a population of cells that are undergoing mitosis, consistent with the PnM cells cycle profile (fig. S6). Using the same approach, we found that the fraction of entirely unpaired tight regions in Slmb and TopII knockdowns increased by 6.80% and 7.50%, relative to mock, and 12.2% and 12.9%, relative to untreated sample (fig. S5A, fig. S10A, B).

### Quantifying genome-wide changes in pairing in knockdowns

In order to quantify the genome-wide changes in pairing observed in knockdowns, we developed a metric that summarized the degree of pairing in each sample, and quantified the genome-wide changes in pairing observed in knockdowns, called aggregated pairing score (APS):

APS = Mode(PS-CS), i.e. the most probable value of PS-CS genome-wide.

We reasoned that APS makes a good estimate for the degree of pairing exhibited by cells genome-wide for two reasons. First, APS has a simple interpretation – since the PS and CS quantify the log_2_ intensity of the main diagonal, APS thus describes the most probable log_2_ ratio of short-distance *thom* contact frequency to short-distance *cis*. Second, APS is stable, since the most probable value in a distribution is not affected by the presence of heavy tails.

For a given pair of conditions, the statistical significance of the difference between their APS was determined with bootstrapping. Specifically, we tested the null hypothesis that the (PS-CS) distributions for both conditions were drawn from the same underlying distribution (i.e. if PS-CS for each condition has a similar distribution). We merged (PS-CS) distributions for both conditions, and then drew 1000X pairs of random samples and calculated the difference of their APS. Finally, we calculated a p-value as a fraction of random sample pairs that had a larger difference of APS than the one observed in our data.

### Eigenvectors

We quantified the compartment structure using of Hi-C maps with eigenvectors of observed/expected Hi-C maps, using a modified procedure from Imakaev *et al*., 2012 (*18*). We performed eigenvector decomposition of the observed/expected 4 kb contacts maps subtracting 1.0 from each pixel. Finally, for each chromosome, we selected the eigenvector showing the highest correlation with the track of the number of genes overlapping each genomic bin. This method for eigenvector detection is available in *cooltools*.

### Other

All custom data analyses were performed in Jupyter Notebooks [Kluyver, Thomas, et al. “Jupyter Notebooks-a publishing format for reproducible computational workflows.” *ELPUB*. 2016.], using matplotlib [Hunter, John D. “Matplotlib: A 2D graphics environment.” Computing in science & engineering 9.3 (2007): 90-95.], numpy [Walt, Stéfan van der, S. Chris Colbert, and Gael Varoquaux. “The NumPy array: a structure for efficient numerical computation.” *Computing in Science & Engineering* 13.2 (2011): 22-30.] and pandas [McKinney, Wes. “pandas: a foundational Python library for data analysis and statistics.” *Python for High Performance and Scientific Computing* (2011): 1-9.] packages. We automated data analyses in command line interface using GNU Parallel [Tange, Ole. “Gnu parallel-the command-line power tool.” The USENIX Magazine 36.1 (2011): 42-47.].

### Detecting chr3R disomy

During the analyses of the WGS and Hi-C reads, we noticed that the maternal homolog (057) of chromosome 3R had consistently higher sequencing coverage than the paternal one (439), from 3.24 Mb and to the telomere. This difference in sequencing coverage was particularly noticeable at the level of the IC balancing weights, which were on average 1.44X higher in this region for the paternal homolog (thus, compensating for lower coverage) comparing to the maternal homolog. Such pattern can be explained by a partial maternal disomy in a fraction of cells. A disomy in a fraction *x* of cells leads to r=(1+x)/(1-x) times higher sequencing coverage of the maternal homolog. Conversely, the observed ratio *r* of sequencing coverage of the two homologs can be explained by partial uniparental disomy in x=(r-1)/(r+1) fraction of cells. Using this approach, we estimate that the chr3R partial disomy is found in 17.9% cells of the untreated Hi-C sample, 20.6% cells of the mock-depleted sample, 21.5% of the Slmb-depleted sample and 19.6% of the TopII-depleted sample.

### RNA-Seq

We mapped the raw RNA-seq data using STAR [https://academic.oup.com/bioinformatics/article/29/1/15/272537] following the same procedure as used by the ENCODE consortium [https://github.com/ENCODE-DCC/long-rna-seq-pipeline/tree/master/dnanexus].

## Supplementary figures

**Fig. S1.**
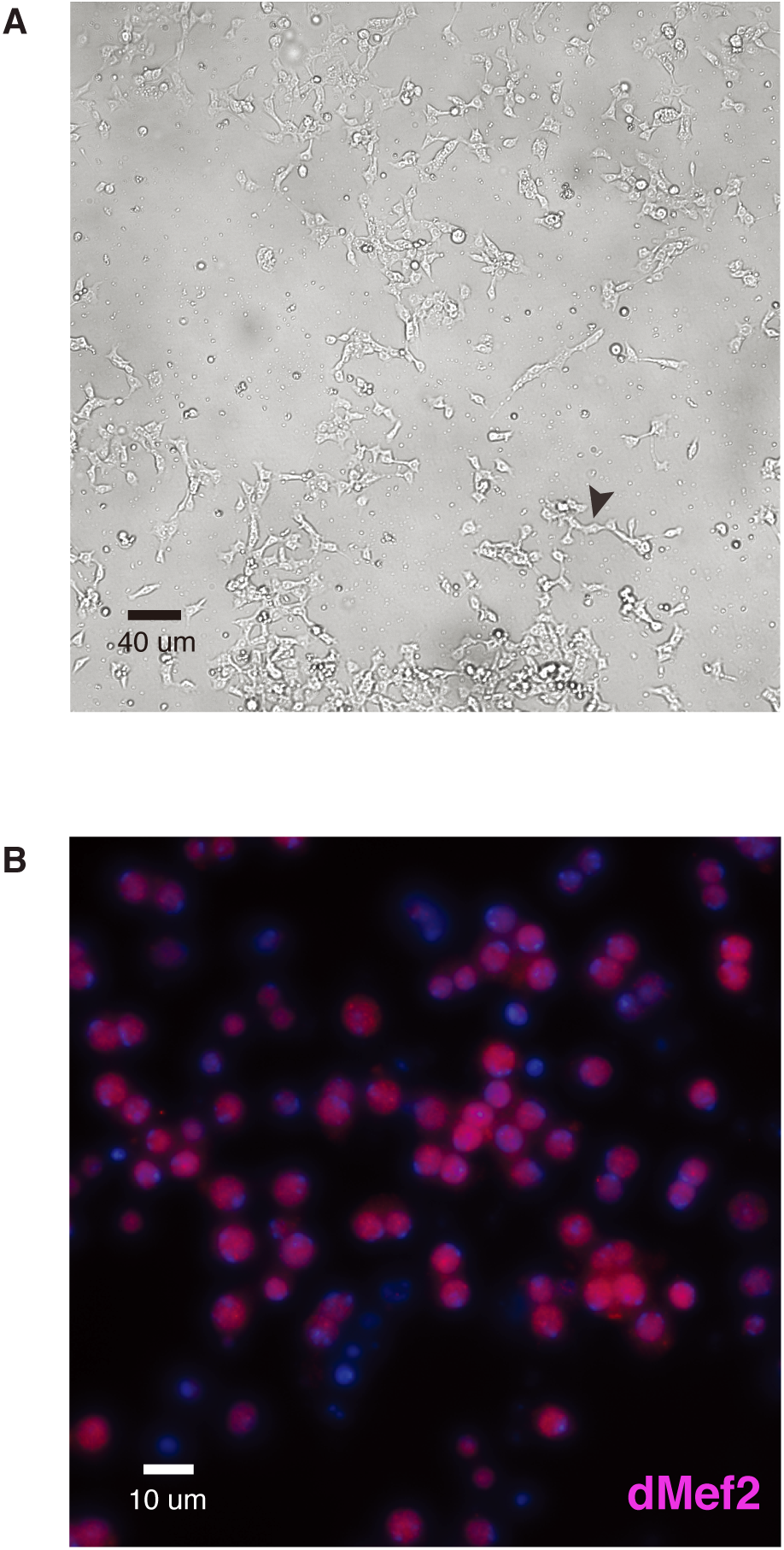
PnM cell line morphology and cell-lineage specification. (A) PnM cell culture consists of adherent, bipolar shaped cells (arrow head). The culture is about 80% confluent, and is ready to passage. Bar= 40 μm (B) PnM cells stained for DAPI and the mesodermal cell marker dMef2, suggested they are predominantly of mesodermal origin. Bar =10 μm.

**Fig. S2.**
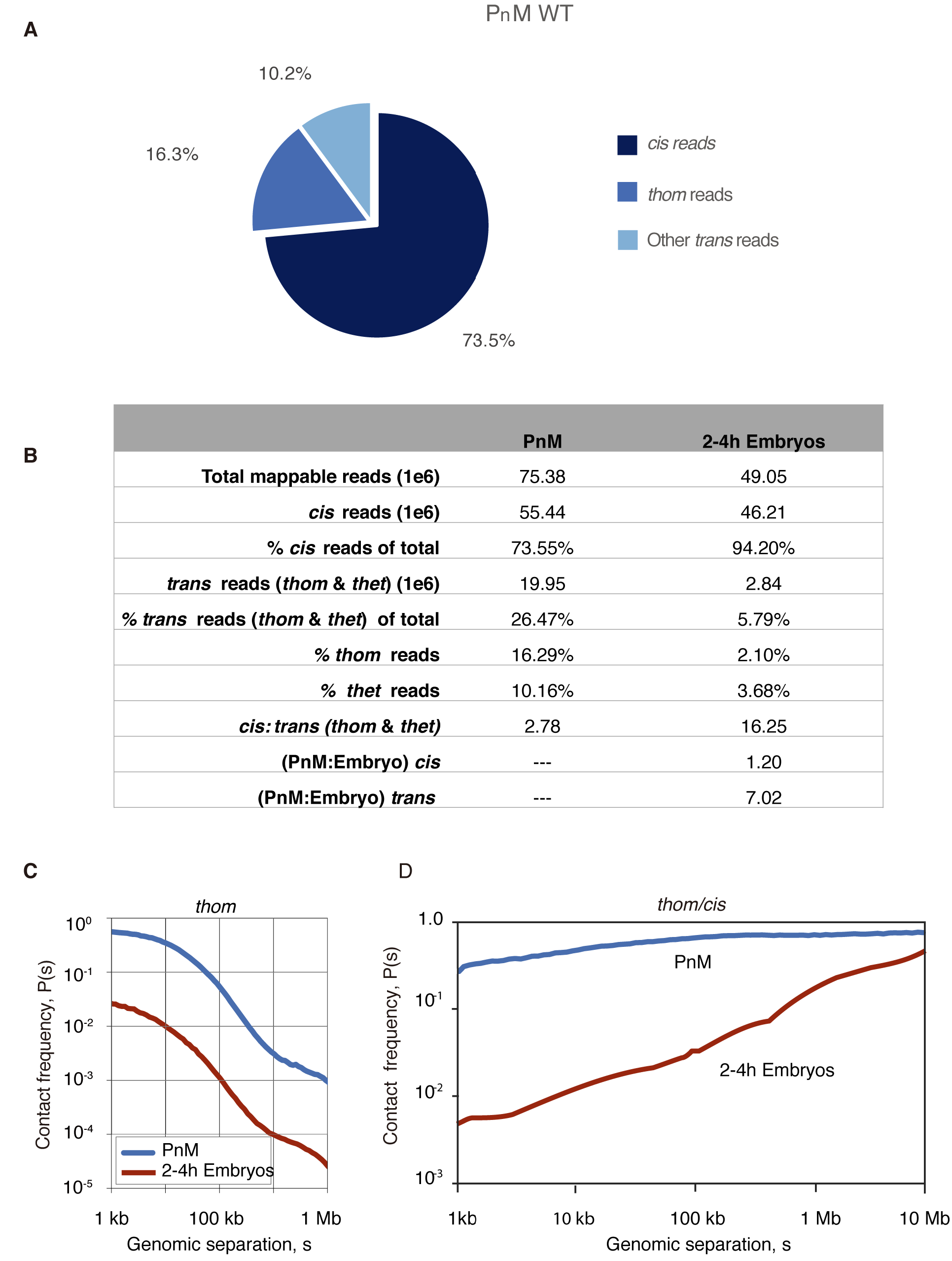
Trans-homolog reads are abundant in PnM cells. (A) Pie chart comparing percentage of *cis* and *trans* reads recovered after haplotyped-mapping to hybrid genome in PnM, where *cis* interactions refer to read pairs with a SNV on each side mapping to the same homolog and *trans* interactions refer to read pair with a SNV on each side mapping to a different homolog or chromosome. Data from PnM untreated cells showing PnM total *trans-*reads are almost a quarter of total reads (B) Comparison of mappable reads in PnM and 2-4 h embryos (Erceg, AlHaj Abed, Golobordko *et al. bioRxiv* (*2*)) shows that the percent of *trans*-homolog (t*hom)* mappable reads in PnM (16.29%) are ~8-fold those of embryos (2.10%) (C) *Thom* signature in PnM cells is greater than *thom* in embryos at all linear separations, s, in kilobases (kb). (D) *Thom*-to-*cis* contact frequencies in 2-4 h embryos and cells. At 100 kb separation in PnM cells, *thom* interaction frequency is almost equal to *cis* interaction frequency. This contrasts to the pairing signature that is observed in 2-4 hr old embryos.

**Fig. S3.**
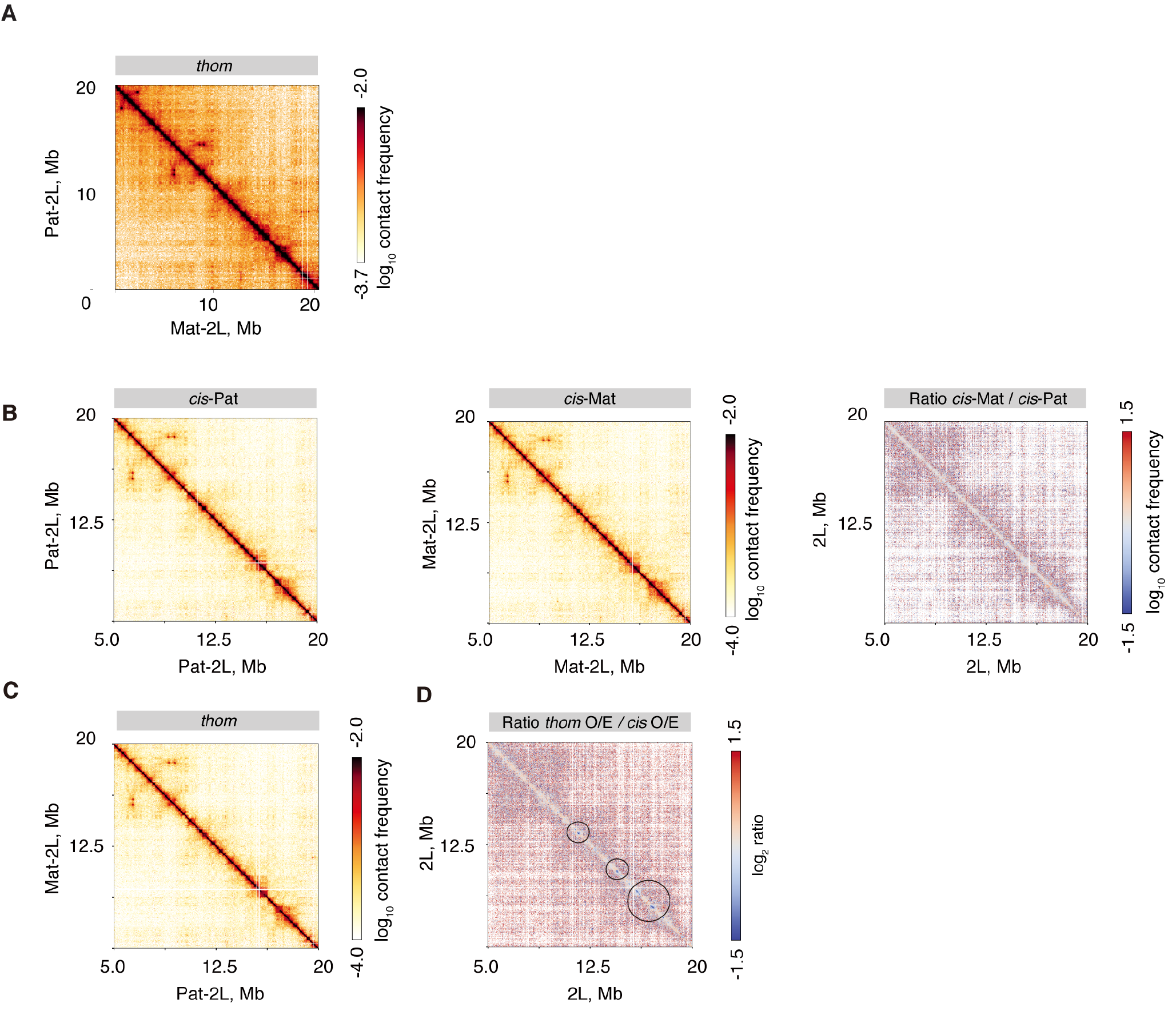
Haplotype maps reveal compartments and concordance of *cis* and *thom* interactions. (A) *Thom* contact map at 2L shows plaid-like pattern representative compartmentalization of homologous chromosomes. (B) Paternal and maternal *cis* contact maps within 5-20 kb at 2L are highly concordant. (C) *Thom* contact map within 5-20 kb at 2L (D) Regions as in B, and C showing *thom* contact frequency (CF) depletion detected in blue (circles pointing to some *thom* depleted regions).

**Fig. S4.**
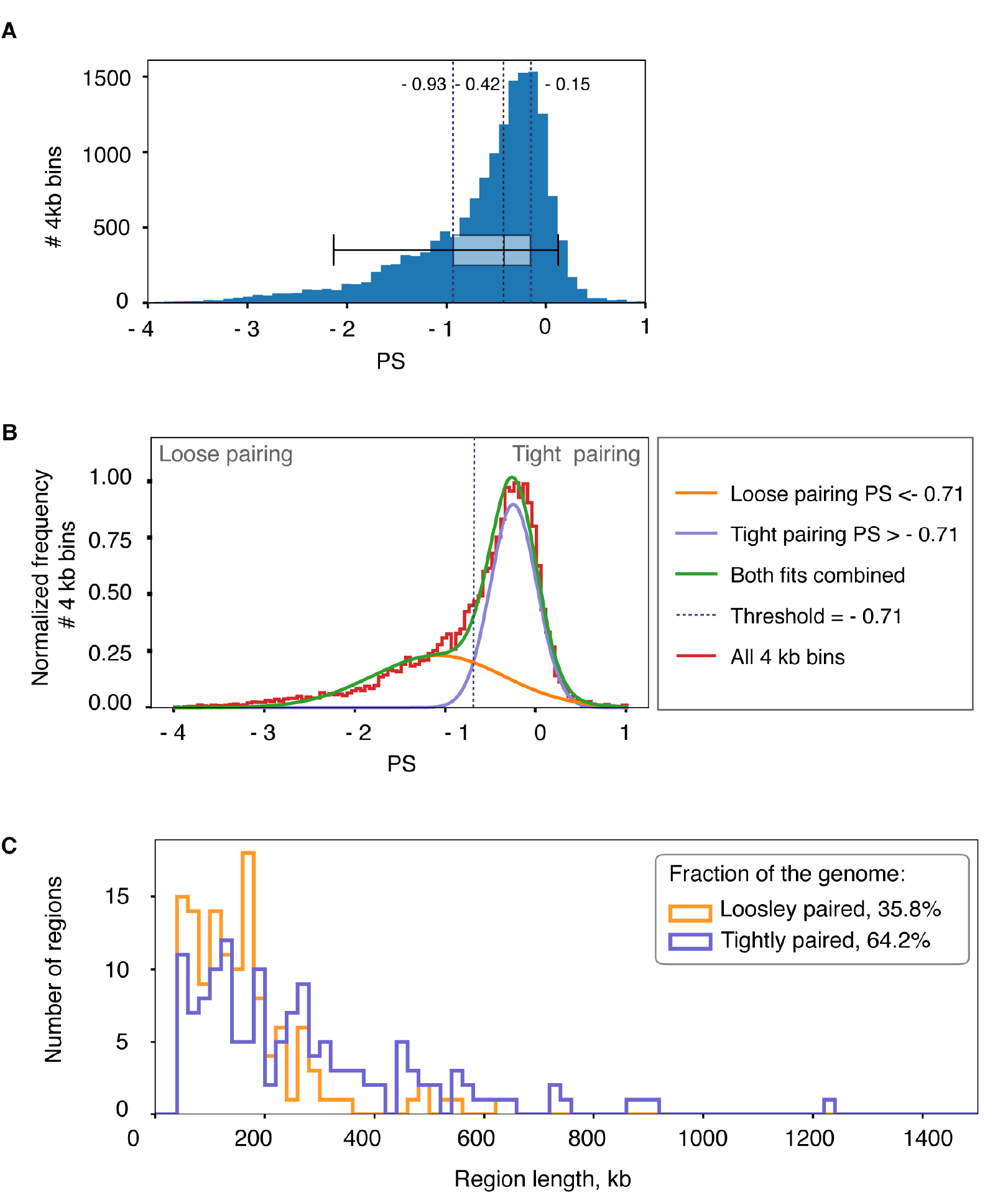
Pairing score distribution and breakdown into tight and loose regions in the genome. (A) Genome-wide distribution of pairing score (PS) in PnM shows a well-pronounced peak at higher values of PS and a tail extending into the lower values of PS. (B) We interpret the PS distribution with a model, where each bin can be either tightly or loosely paired. We can separate the peak from the tail of the distribution by fitting it with a sum of two Gaussians. All genomic bins with PS < −0.71 are considered loosely paired (since they are more likely to belong to the low-PS Gaussian) and the bins with PS > −0.71 are considered tightly paired. (C) The length distribution of loosely and tightly paired regions

**Fig. S5.**
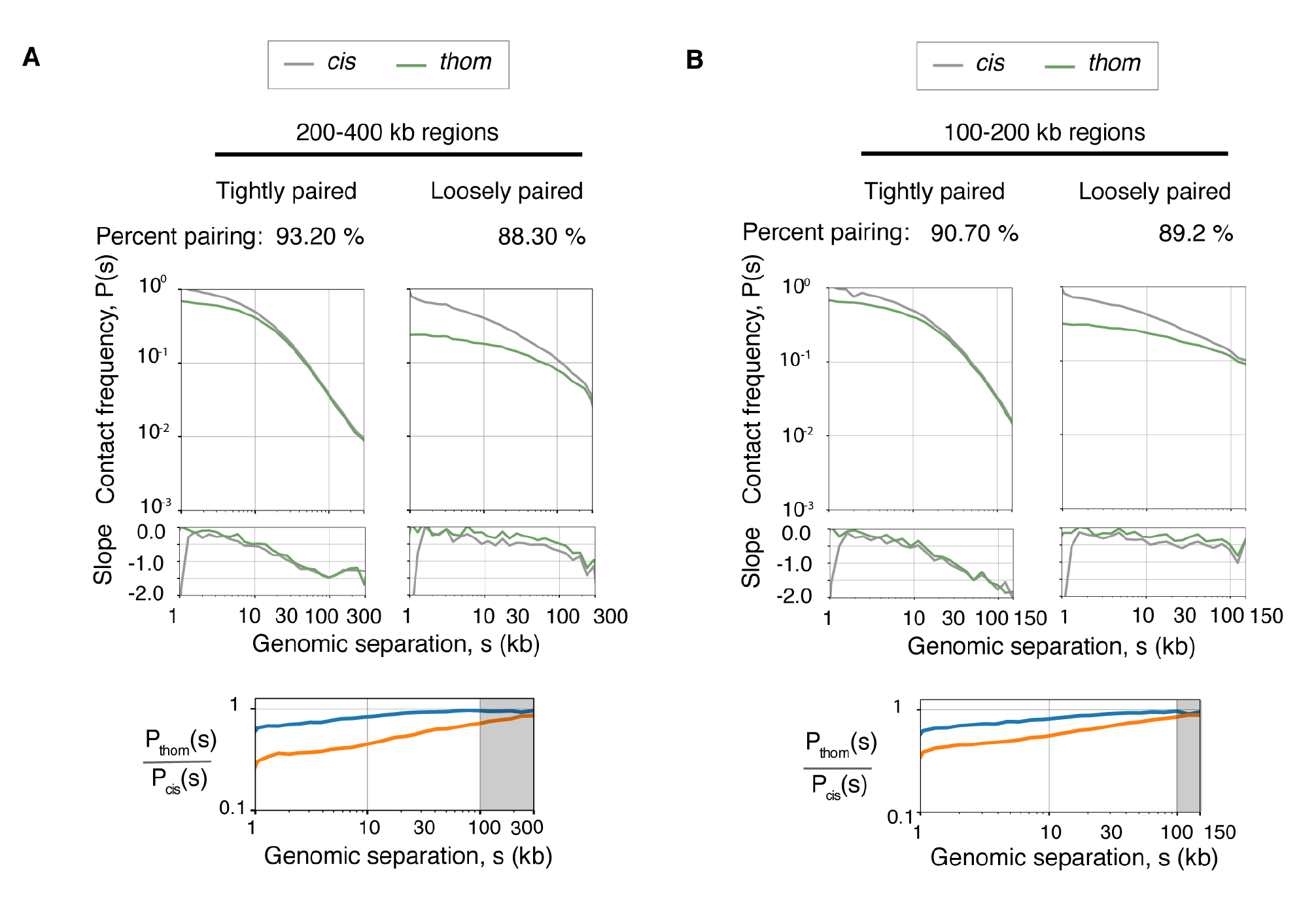
P_thom_(s), P_cis_(s) for tight and loose regions of different sizes. Top, P_thom_(s), P_cis_(s). Middle, slope, and bottom, is *thom/cis* CF within tightly and loosely paired regions. (A) 200-400 kb length and (B) 100-200 kb length. In tighly paired regions, *thom* and *cis* CF curves show two modes of decay, shallow and steep, while in loosely paired regions, the curves have only one, shallow mode. Shaded region, is the percent pairing determined as the geometric mean of the *thom/cis* ratio at longer genomic separations (100-300 kb, or 100-150 kb).

**Fig. S6.**
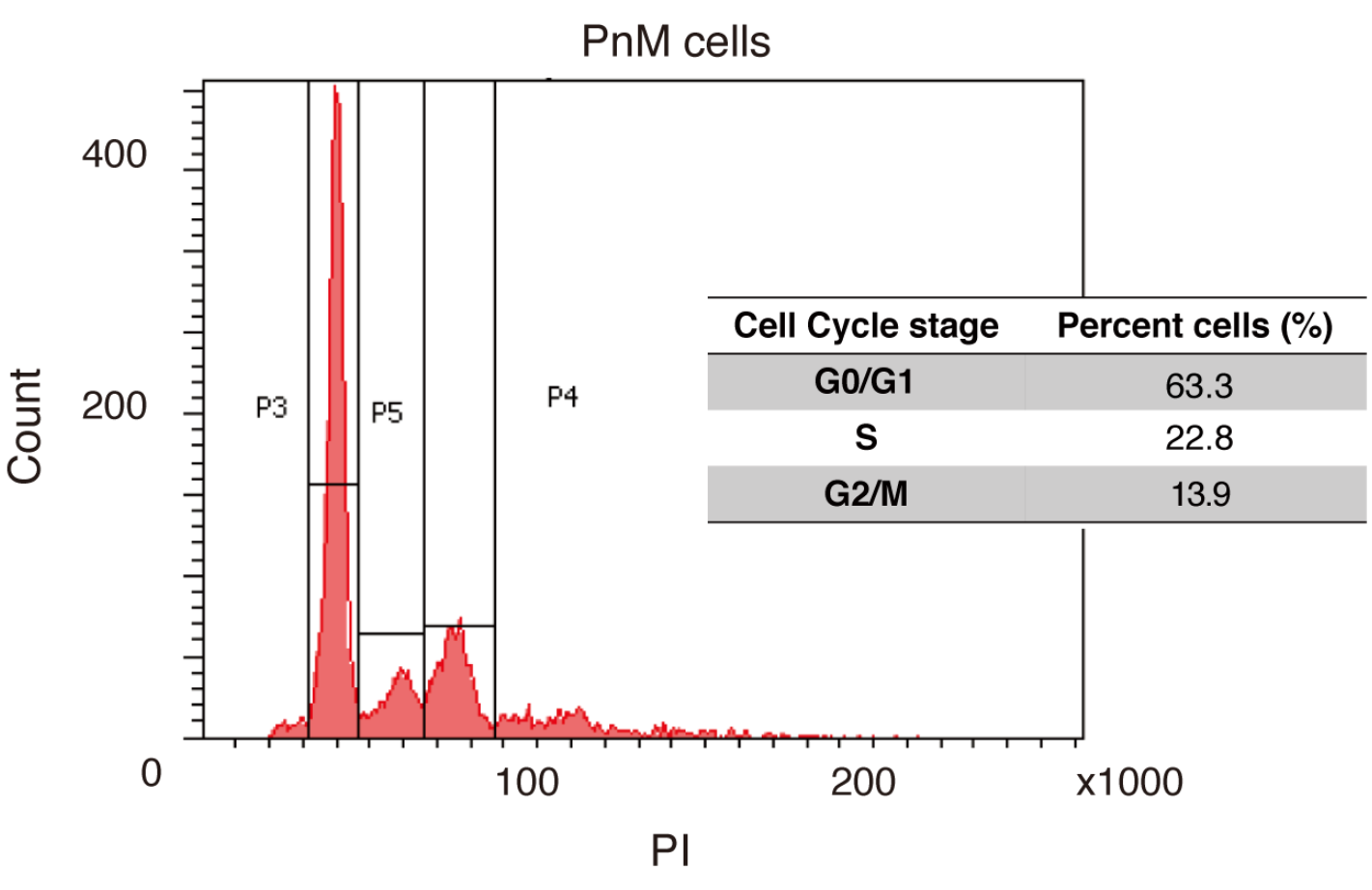
Cell cycle analysis of PnM cells. Staining of cells with propidium Iodide at 70% confluency shows that they are predominantly in G1.

**Fig. S7.**
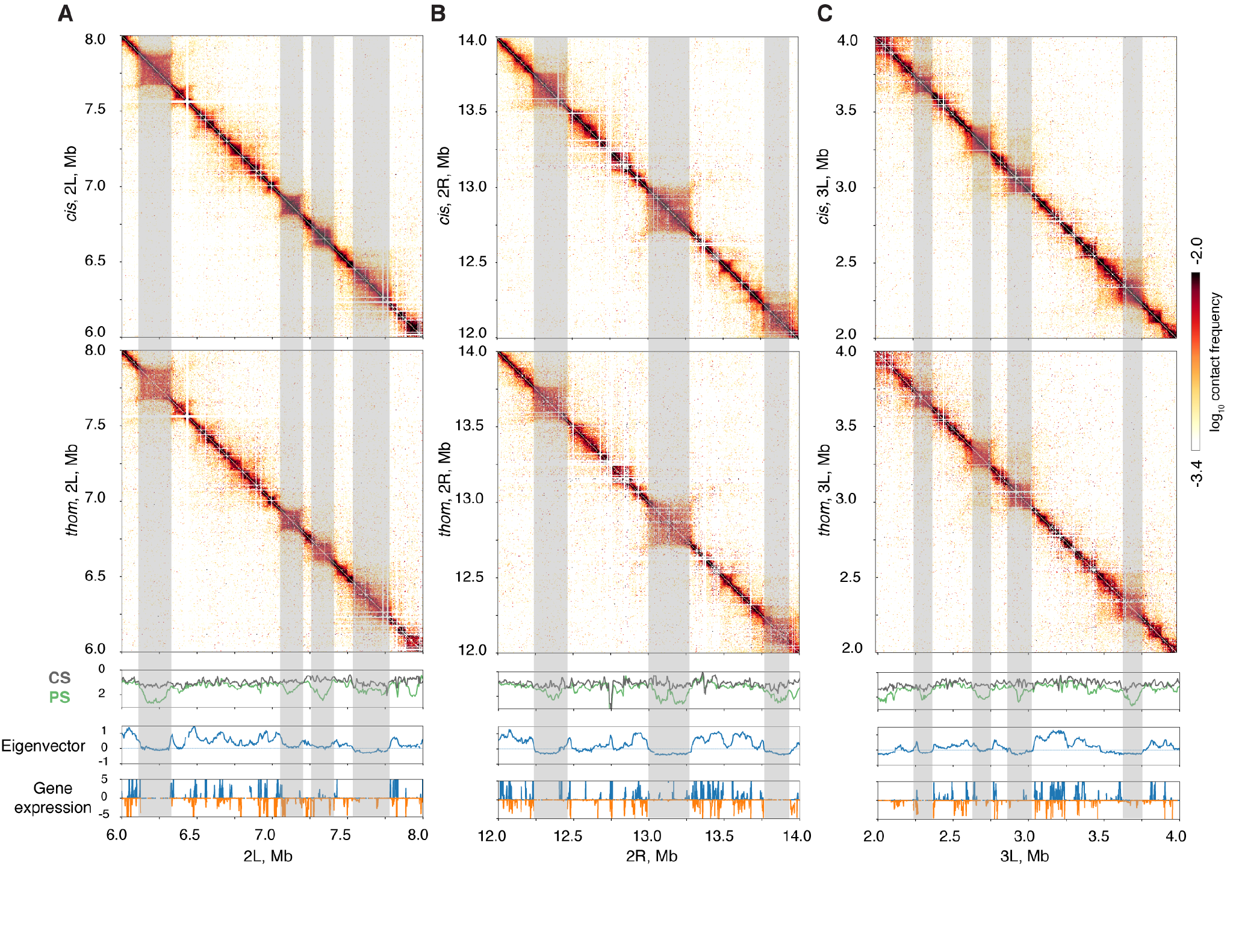
Representative examples of pairing relative to compartments and gene expression. (A-C) 2 Mb-long example regions described with *cis* and *thom* contact maps (two upper panels), pairing (PS) and *cis* scores (CS) (middle panel) and measures of transcriptional activity, eigenvector and RNA-seq (two lower panels). High PS correlates with high levels of gene expression, and enrichment of A compartments. Some weakly paired regions are shaded, indicating a lower eigenvector rank, and generally lower expression levels. (A) 2L, 6-8 Mb, (B) 2R, 12-14 Mb and (C) 3L, 2-4 Mb.

**Fig. S8.**
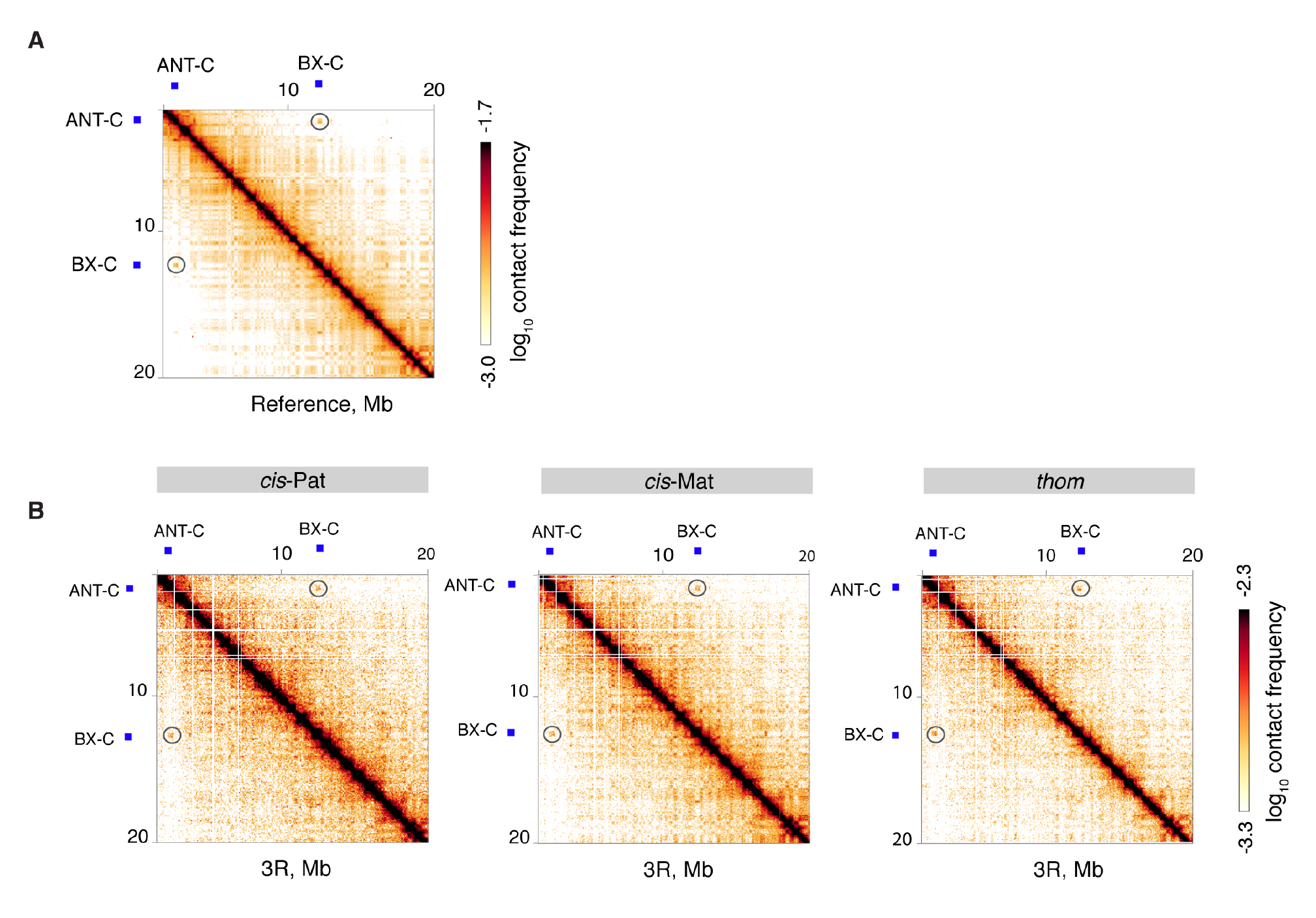
*Thom* interaction of ANT-C and BX-C. (A) Long-range *cis*-interactions that were detected in previous studies between ANT-C and BX-C (*19*) are detected in PnM reference map (non-allele-specific mapping), (B) and *thom* and *cis* maps, indicating that this well-known interaction is within a paired compartment in our haplotype-maps.

**Fig. S9.**
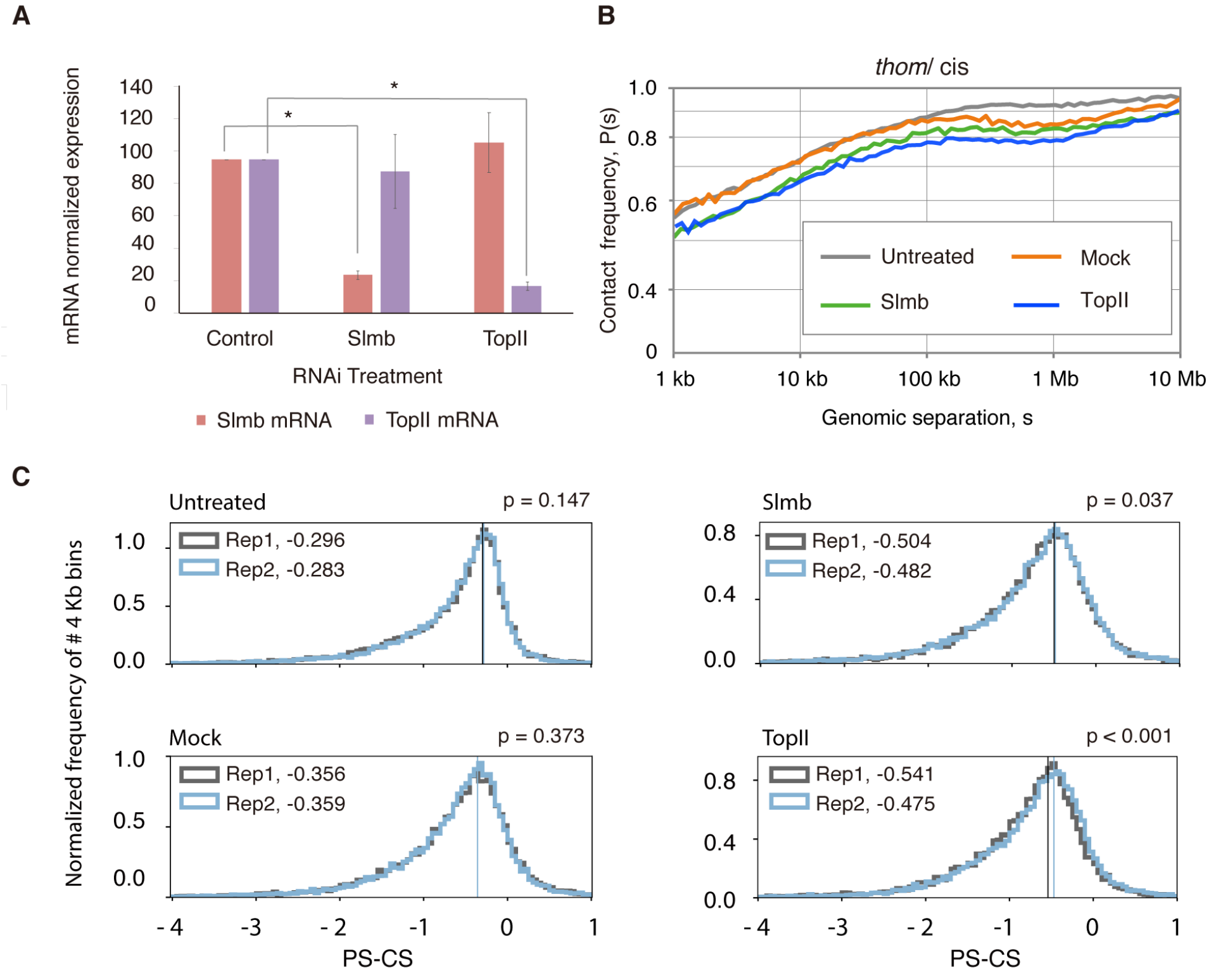
The effect of knocking down Slmb or TopII. (A) Quantitative PCR confirmed efficient knockdown of Slmb and TopII. Their expression levels are normalized to Act5c and RP49 expression after RNAi in PnM cells. There is a significant drop in the levels of Slmb and TopII mRNA compared to the control mock treatment. Asterisks denote a significant reduction from control (P < 0.0001, unpaired t-test). S.d are shown for at least 3 biological replicates. (B) The ratio of *thom*-to-*cis* CF as a function of genomic separation, s, in kilobases (kb), in Slmb, and TopII RNAi sample show a drop in *thom*-to-*cis* frequency compared to mock at distances below 0.1 Mb and at all genomic separations compared to the untreated PnM cells (C) *Thom* contact frequency is reduced in Slmb and TopII RNAi samples compared to untreated cells, and mock replicates, for two biological replicates as illustrated by a drop in the PS-CS distribution. P-values determined using bootstrapping (Supplementary Material).

**Fig. S10.**
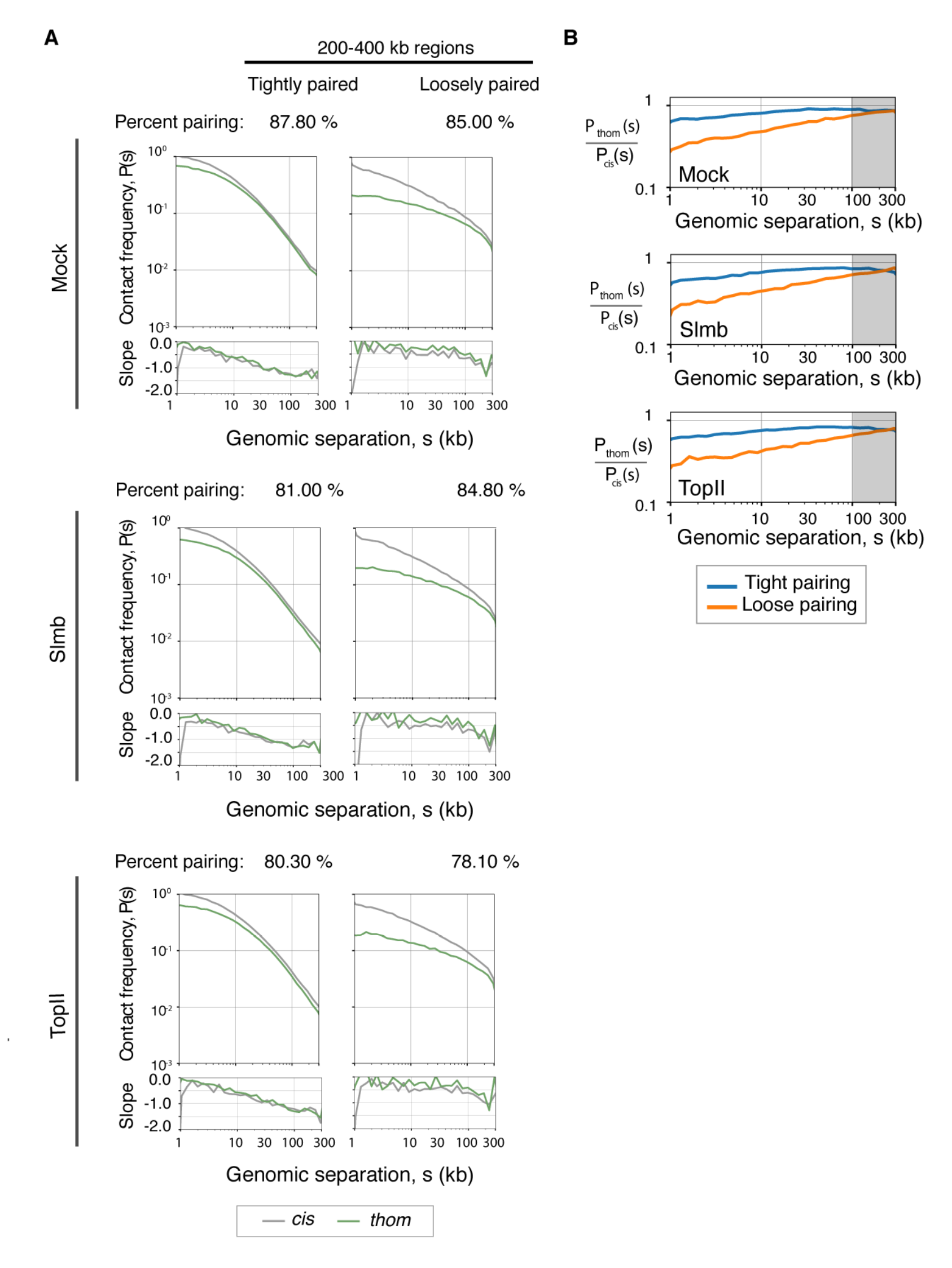
The effect of knocking down Slmb or TopII on 200-400 kb regions of tight and loose pairing. (A) Plots of P(s)_thom_ and and P(s)_cis_ slope within tightly (left) and loosely paired (right) regions of 200-400 kb length. (B) The P(s)_thom_ */* P(s)_cis_ ratio relative to genomic separation, s, in tightly and loosely paired regions. Shaded region, shows that the percent pairing in (A) is determined as the geometric mean of the *thom/cis* ratio at longer genomic separations (100-300kb). A comparison between tight and loose pairing *thom* and *cis* CF shows a drop in both tight and loose pairing for Slmb and TopII relative to untreated cells.

## Supplementary Tables

**Table S1.** Summary of SNV frequency per chromosome

| Chrom | Chrom size | Indels |  |  | SNVs |  |  | het SNVs/kb |
| --- | --- | --- | --- | --- | --- | --- | --- | --- |
|  |  | #057 | #439 | hom | #057 | #439 | hom |  |
| chr2L | 23011544 | 5820 | 5910 | 9415 | 66894 | 70989 | 68091 | 5.99 |
| chr2R | 21146708 | 5144 | 5396 | 8828 | 55659 | 58689 | 60606 | 5.41 |
| chr3L | 24543557 | 6068 | 5599 | 10374 | 67299 | 64261 | 73011 | 5.36 |
| chr3R | 27905053 | 6494 | 6402 | 9676 | 66777 | 68709 | 65627 | 4.86 |
| chr4 | 1351857 | 1 | 4 | 136 | 2 | 15 | 929 | 0.01 |
| chrX | 22422827 | 8405 | 8386 | 5193 | 40674 | 44055 | 38063 | 3.78 |

**Table S2.**
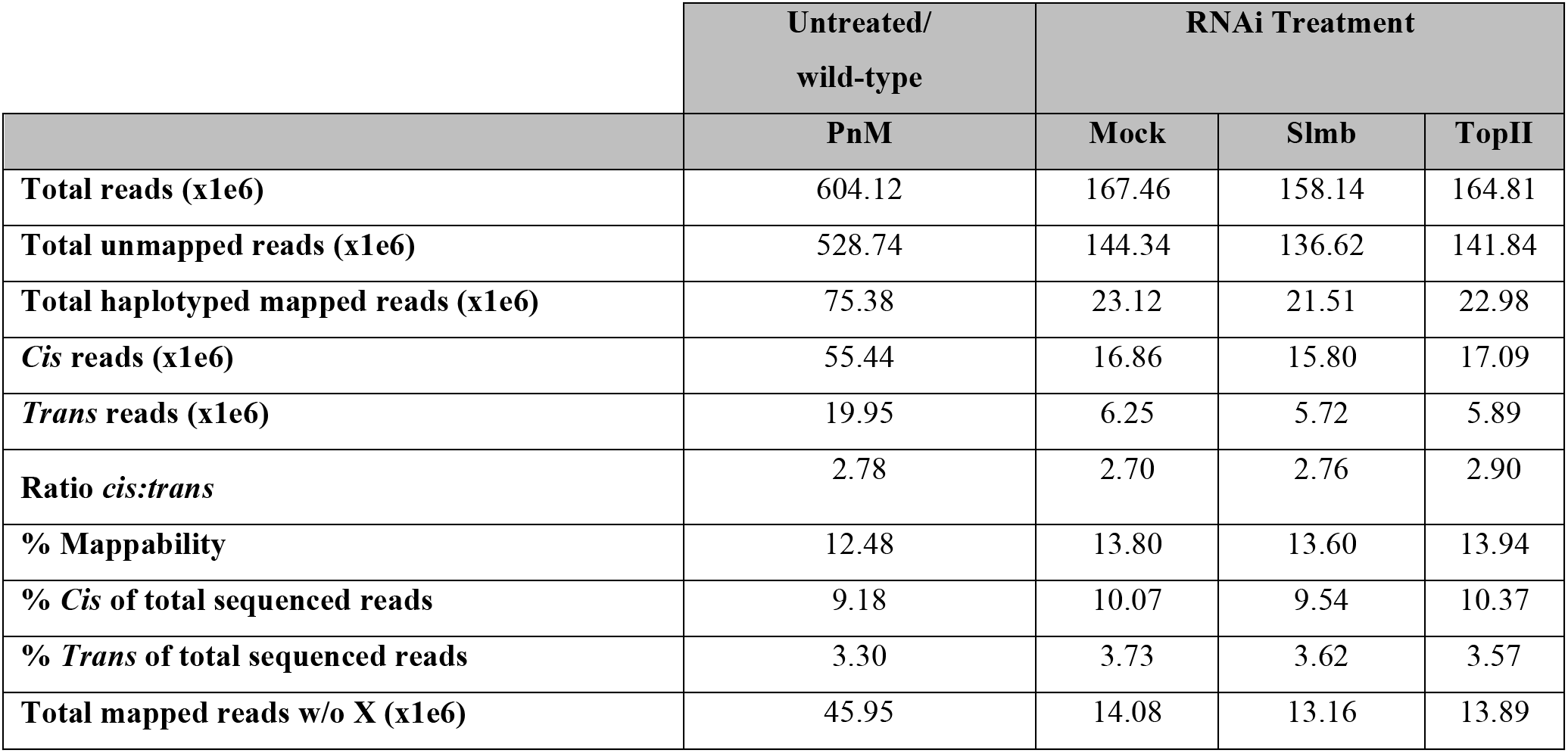
Summary of haplotyped read pairs used in Hi-C haplotyped-mapping for wild-type PnM and RNAi treated cells, with two biological replicates merged per sample.

|  | Untreated/<br>wild-type | RNAi Treatment |  |  |
| --- | --- | --- | --- | --- |
|  | PnM | Mock | Slmb | TopII |
| **Total reads (x1e6)** | 604.12 | 167.46 | 158.14 | 164.81 |
| **Total unmapped reads (x1e6)** | 528.74 | 144.34 | 136.62 | 141.84 |
| **Total haplotyped mapped reads (x1e6)** | 75.38 | 23.12 | 21.51 | 22.98 |
| ***Cis* reads (x1e6)** | 55.44 | 16.86 | 15.80 | 17.09 |
| ***Trans* reads (x1e6)** | 19.95 | 6.25 | 5.72 | 5.89 |
| **Ratio *cis:trans*** | 2.78 | 2.70 | 2.76 | 2.90 |
| **% Mappability** | 12.48 | 13.80 | 13.60 | 13.94 |
| **% *Cis* of total sequenced reads** | 9.18 | 10.07 | 9.54 | 10.37 |
| **% *Trans* of total sequenced reads** | 3.30 | 3.73 | 3.62 | 3.57 |
| **Total mapped reads w/o X (x1e6)** | 45.95 | 14.08 | 13.16 | 13.89 |

**Table S3.**
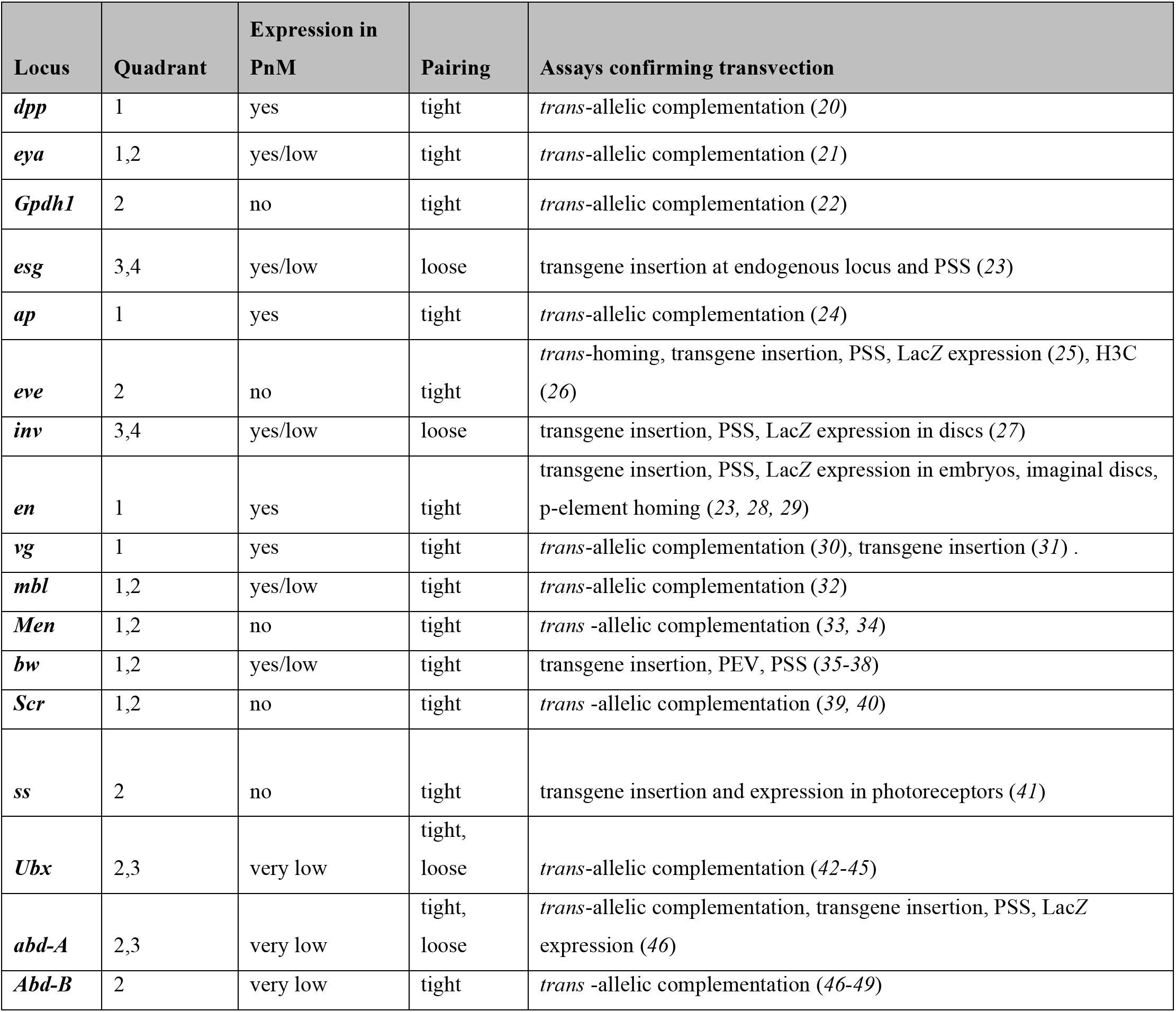
Examples of known transvected loci on the autosomes and their location in Fig. 3D. Quadrant 1, and 2, represents regions in the genome that are tightly paired and are expressed or not expressed at all *Pairing sensitive silencing (PSS), high resolution chromosome conformation capture (H3C), position effect variegation (PEV). *The summary of loci above include at least one endogenous locus study, or endogenous locus regulatory region*.

**Table S4.** Summary of different pairing assays for Slmb and TopII knockdown samples relative to mock sample.

| Assay | Slmb | TopII |
| --- | --- | --- |
| qPCR mRNA depletion vs. mock | 75.2 % | 82.5 % |
| Average FISH drop in colocalization vs. mock | 12.1 % | 10.7 % |
| Average FISH drop in colocalization vs. untreated sample | 22.2 % | 20.8 % |
| CPS change vs. mock | 7.5 % | 7.1 % |
| CPS change vs. untreated sample | 11.0 % | 10.6 % |

**Table S5.** Oligopaints probe target library summary

| Probe target | dm3 Coordinates | OligoLibraryID | Total # probes | Target size (Mb) | Secondary oligo ID used |
| --- | --- | --- | --- | --- | --- |
| **28B** | chr2L: 7,256,488-7,936,487 | Mycroarray-6151 | 10000 | 0.680 | Sec5 |
| **69C** | chr3L: 12,170,682-12,844,681 | Mycroarray-6150 | 10000 | 0.674 | Sec1 |
| **BX-C** | chr3R: 12,482,502-12,797,965 | Mycroarray-6010 | 2394 | 0.316 | Sec6 |
| **16E** | chrX: 17,406,557-18,106,556 | Mycroarray-6148 | 10000 | 0.700 | Sec6 |
| **HOPs 2L** | chr2L:10,731-1,999,862 | Mycroarray | 2410 | 2.000 | 2L-057: Sec6, 2L-439: Sec5 |
| **HOPs 3L** | chr3L: 19,809-1,999,387 | Mycroarray | 2304 | 2.000 | 3L-057: Sec6, 3L-439: Sec5 |

**Table S6.**
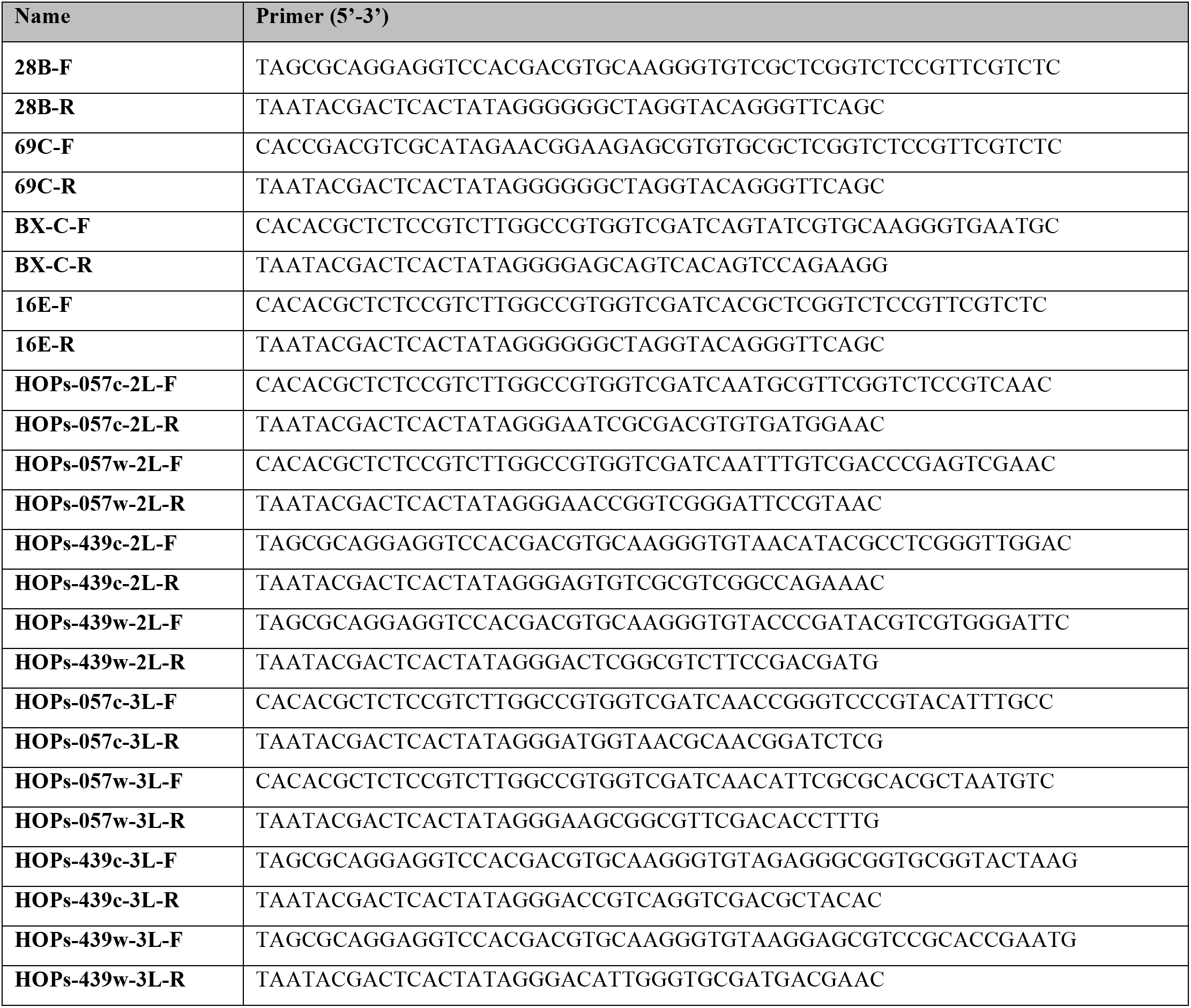
Oligopaints libraries PCR primers

**Table. S7.**
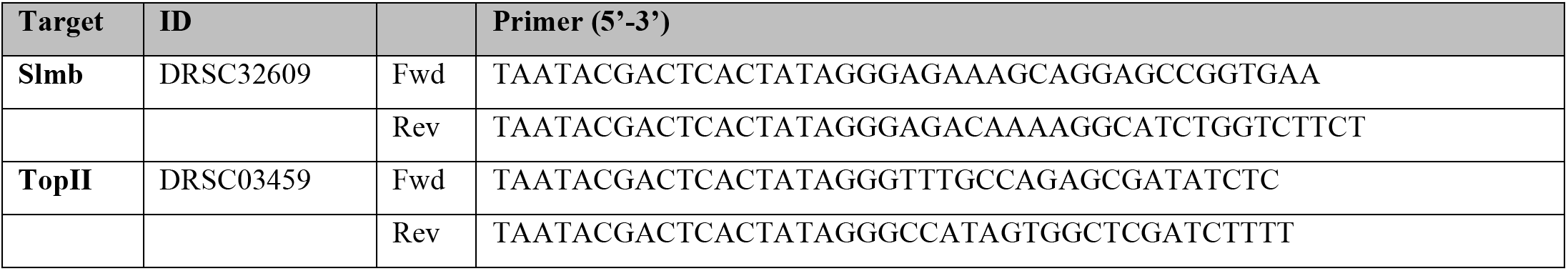
dsRNA synthesis primers

**Table. S8.**
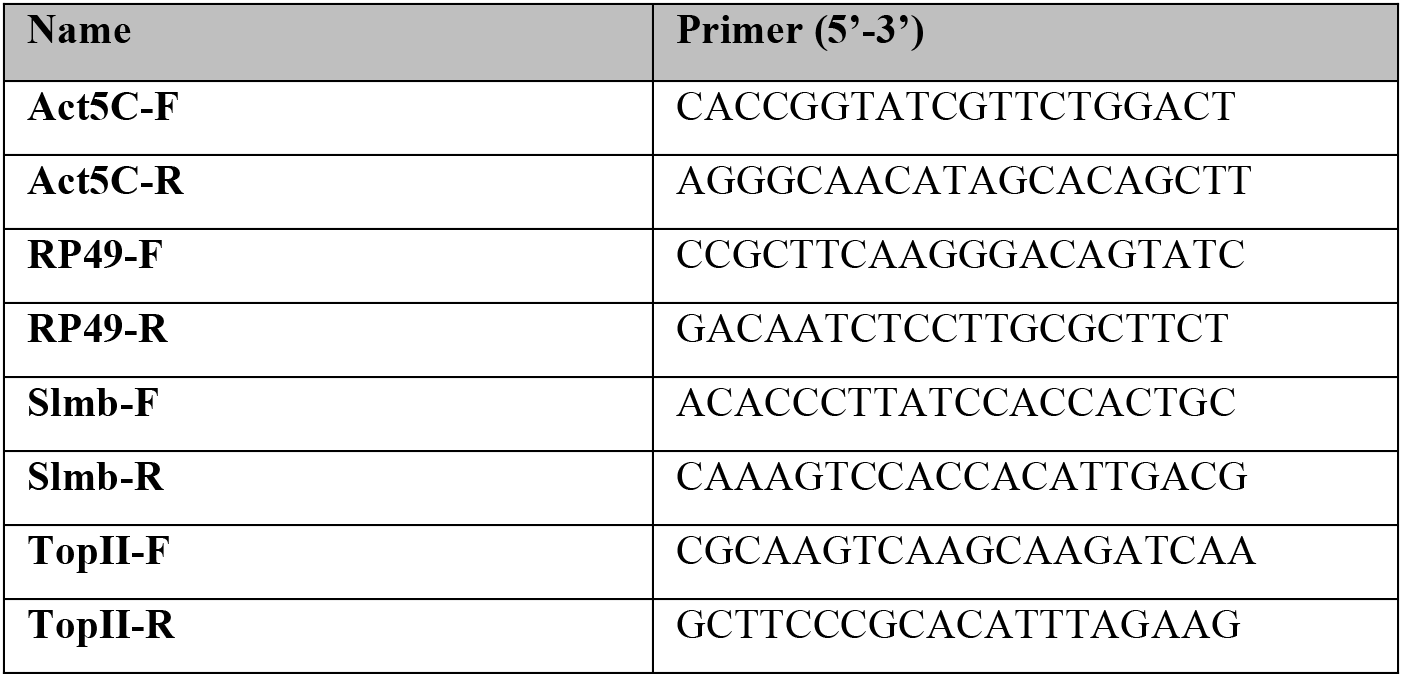
qPCR primers

